# High-precision spatial analysis of mouse courtship vocalization behavior reveals sex and strain differences

**DOI:** 10.1101/2021.10.22.464496

**Authors:** G. Oliveira-Stahl, S. Farboud, M. L. Sterling, J. J. Heckman, B. van Raalte, D. Lenferink, A. van der Stam, C. J. L. M. Smeets, S. E. Fisher, B. Englitz

**Affiliations:** Department of Neurophysiology, Donders Institute for Brain, Cognition and Behaviour, Radboud University, Nijmegen, The Netherlands; Language and Genetics Department, Max Planck Institute for Psycholinguistics, Nijmegen, The Netherlands; Donders Institute for Brain, Cognition and Behaviour, Radboud University, Nijmegen, The Netherlands

**Keywords:** mouse social interaction, ultrasonic vocalizations, USV, vocal communication, sound localization, heterozygous Foxp2-R552H mice

## Abstract

Mice display a wide repertoire of vocalizations that varies with sex, strain, and context. Especially during social interaction, mice emit sequences of ultrasonic vocalizations (USVs) of high complexity. As animals of both sexes vocalize, a reliable attribution of USVs to their emitter is essential.

The state-of-the-art in sound localization for USVs in 2D allows spatial localization at a resolution of multiple centimeters. However, animals interact at closer ranges, e.g. snout-to-snout. Hence, improved algorithms are required to reliably assign USVs. We present a novel algorithm, SLIM (Sound Localization via Intersecting Manifolds), that achieves a 3-fold improvement in accuracy (12-14.3mm) using only 4 microphones and extends to many microphones and localization in 3D. This accuracy allows reliable assignment of 84.3% of all USVs in our dataset.

We apply SLIM to courtship interactions between adult C57Bl/6J wildtype mice and those carrying a heterozygous Foxp2 variant (R552H). The improved spatial accuracy reveals detailed vocalization preferences for specific spatial relations between the mice. Specifically, vocalization probability, duration, Wiener entropy, and frequency level differed in particular spatial relations between WT females, Foxp2-R552H and WT males.

In conclusion, the improved attribution of vocalizations to their emitters provides a foundation for better understanding social vocal behaviors.

## Introduction

Mice emit ultrasonic vocalizations (USVs) during a variety of behaviors. For instance, when a pup is isolated from the nest, it exclaims a distress or isolation call to warn its mother, sometimes dramatically referred to as the *whistle of loneliness* (Zippelius and Schleidt, 1956). In contrast, adult animals vocalize predominantly in the presence of other mice to mediate essential social behaviors, such as territorial disputes and courtship. The USVs of mice differ depending on their age, genetic background, sex, and behavioral state (Heckman et al., 2016). Vocalization sequences produced during courtship have been described as complex and non-random (Holy and Guo, 2005), suggesting a potential conveyance of information. Accordingly, behavior in animals hearing the vocalizations (’receiver animals’) is also susceptible to these changes as can be seen in playback studies which have shown that mice prefer different types of vocalizations (Chabout et al., 2015; Hammerschmidt et al., 2009; Liu and Schreiner, 2007).

Recent interest in the genomic contributions to human speech development and associated disorders, in particular the influence of genes such as *FOXP2* (Lai et al., 2001), has broadened the possibilities for studying vocal behaviors in mice. A variety of *Foxp2* mouse lines have been developed (French and Fisher, 2014) that carry mutations matching those found in human cases of speech/language disorder (Fujita-Jimbo and Momoi, 2014; Fujita et al., 2008; Gaub et al., 2010; Groszer et al., 2008), as well as knock-out variants (Shu et al., 2005) and partially humanized *Foxp2* lines (Enard et al., 2009; Hammerschmidt et al., 2015).

Over the last few years, the study of vocal interactions between mice has been advanced by technical improvements in sound source localization (Heckman et al., 2017; Neunuebel et al., 2015; Sangiamo et al., 2020; Warren et al., 2018). Previously, we developed an algorithm that improved the spatial precision of USV localization >3-fold in a single dimension (Heckman et al., 2017), allowing more accurate USV attribution during close-range dyadic interactions (e.g. face-to-face). While we reported a distribution of the assigned USVs to male and female animals that was similar to Neunuebel *et al*. (Neunuebel et al., 2015), we found that the multiple basic properties of USVs differed between sexes. Further progress to understand vocal behavior during dyadic interactions can be made by quantifying the animals’ behavior in greater detail (Bobrov et al., 2014; Rao et al., 2014; Wolfe et al., 2011).

In the present study, we generalize our USV localization algorithm ’SLIM’ (Sound Localization via Intersecting Manifolds) from one to 2D or multiple dimensions in order to assess USVs during social courtship interactions such as facial touch, anogenital sniffing, and chasing. We achieve an average localization error of 14.3 mm for all USVs, 13.1 mm when selecting for a subset of reliably assigned USVs constituting 84.3% of the total set, and 11.9 mm for cases in which the animals are widely separated from each other (>100 mm). The present accuracy constitutes a >3-fold improvement over the previously reported accuracies (Neunuebel et al., 2015).

We utilize SLIM-based localization to study vocalization behavior in male-female courtship interactions for wildtype (WT) females with males that were either WT or carried a heterozygous etiological Foxp2 mutation (Foxp2-R552H), all three on a C57Bl/6J background (Groszer et al., 2008), as well as CBA/CaJ WT females and males. Vocalizations of Foxp2-R552H mice differ from male WT mice in duration, Wiener entropy, and level. Further, we find differences in vocalization probability, duration, frequency range, and Wiener entropy for particular spatial relations between male and female mice. In summary, the present study lays the foundation for a more advanced understanding of vocal interactions through improved attribution of vocalizations to their emitters.

## Results

We developed a novel technique for localizing ultrasonic vocalizations (USVs) which enabled a refined analysis of the vocalization behavior of mice during social interaction. We analyzed the properties of USVs and the relative positions of emitter and receiver for both sexes and different strains (in total N=40 mice, 170 recordings with USVs, 38,092 USVs, see Table 1 for details). In the following we compared (1) male C57Bl/6J WT vs female C57Bl/6J WT mice during social interaction and (2) male Foxp2-R552H vs littermate controls (C57Bl/6J WT) both interacting with C57Bl/6J WT females. As a methods control, we compared the (3) accuracy of localization using SLIM on 3 (all CBA/CaJ WT USVs) vs 4 microphones (all Foxp2-R552H and C57Bl/6J WT USVs). Comparison between CBA/CaJ WT and the other groups was not performed, as the accuracy of localization differed and frame-by-frame tracking of CBA/CaJ WT mice was not available due to the lack of identifying markers.

**Table 1:**
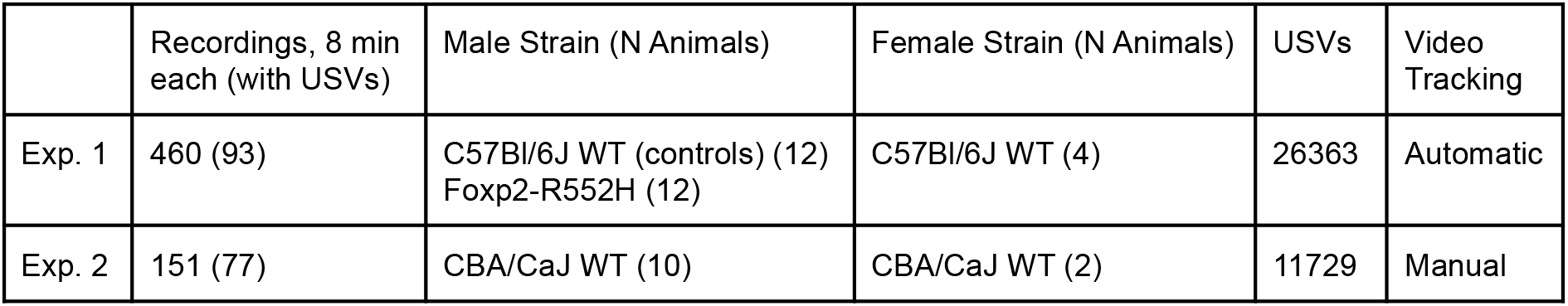
Overview of the properties of the two experiments conducted, regarding the strain differences, number of recordings, type of video track and collected USVs.

|  | Recordings, 8 min each (with USVs) | Male Strain (N Animals) | Female Strain (N Animals) | USVs | Video Tracking |
| --- | --- | --- | --- | --- | --- |
| Exp. 1 | 460 (93) | C57Bl/6J WT (controls) (12)<br>Foxp2-R552H (12) | C57Bl/6J WT (4) | 26363 | Automatic |
| Exp. 2 | 151 (77) | CBA/CaJ WT (10) | CBA/CaJ WT (2) | 11729 | Manual |

A male and a female mouse interacted freely on an elevated platform inside a soundproof booth (Fig. 1A) while their vocalizations were recorded using multiple (3 or 4, see Methods) ultrasonic microphones, and their movements were recorded from above using a high-speed camera (Fig. 1B, red: female; blue: male). Mice vocalized frequently during these social encounters (instantaneous rates typically 8-10 USVs/s, Fig. 1C), in particular during close interaction (Fig. 1D), necessitating high-precision techniques for localizing USVs in space and attributing them to one of the interacting mice.

**Figure 1:**
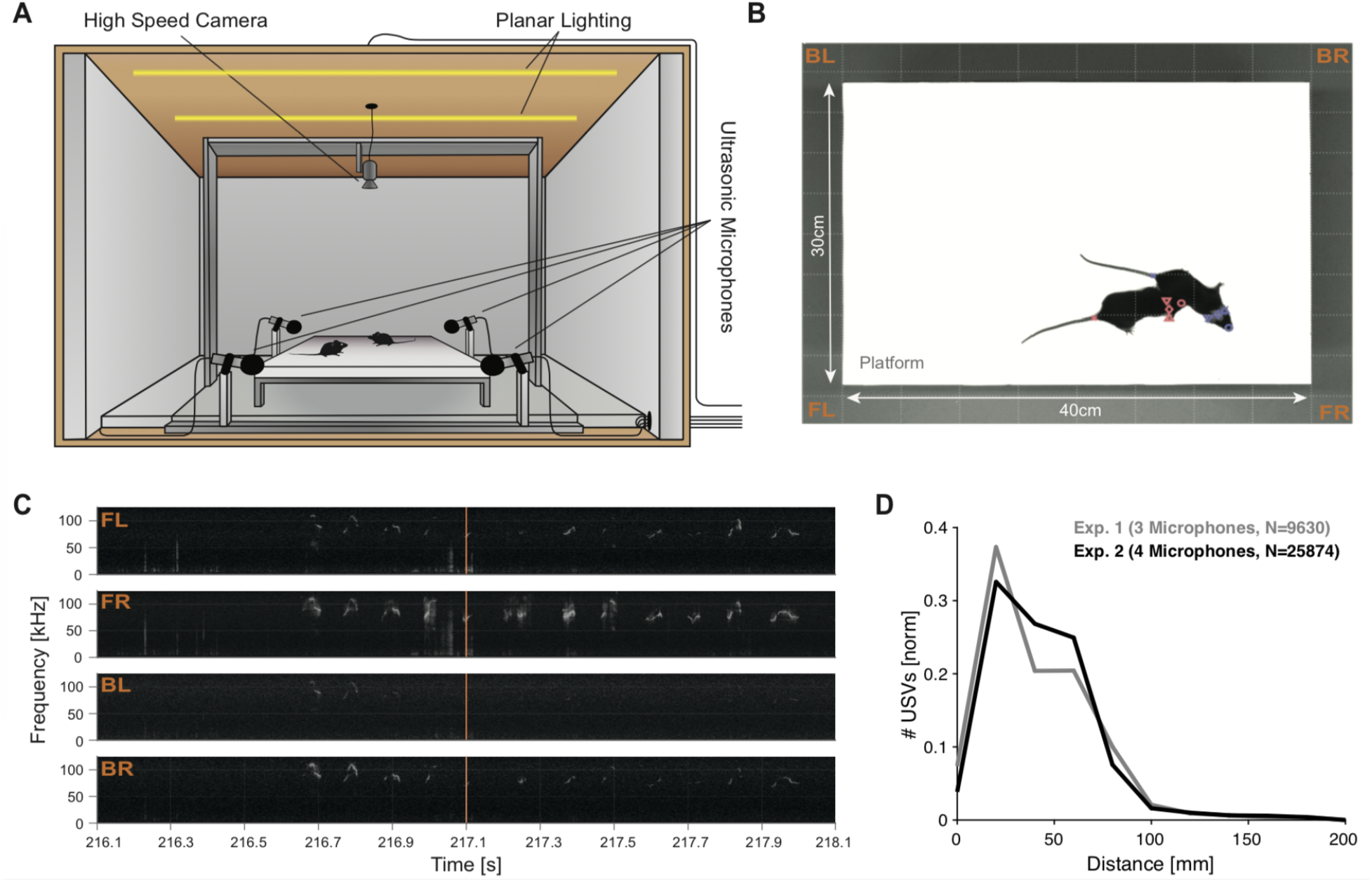
The study of mouse vocalizations during natural behavior requires attribution of individual vocalizations to individual mice. **A** A pair of female and male mice interacted freely on an elevated platform. Their spatial location, behavior and vocal production was monitored with a high-speed camera (placed directly above) and 4 ultrasonic microphones surrounding the platform. The whole setup was situated inside a sound-proof, ultrasonically anechoic box which was uniformly illuminated using a planar array of LEDs. **B** Animals were easily distinguishable against the white, anechoic platform. **C** Vocalizations occurred frequently during most experiments, in particular during social interaction of the animals (Frame in **B** at time = 217.1s). **D** In the present paradigm, the majority of vocalizations were emitted when the animals were in rather close proximity (black: 4 microphone setup, gray: 3 microphone setup), typically below 10 cm snout-to-snout distance, shown on the abscissa.

We automatically tracked both animals on the platform in 2D using DeepLabCut (Mathis et al., 2018) and subsequent post-processing (Fig. 2A, red: female; blue: male; all tracking points shown for both animals). The resulting behavioral tracks were accurate to a few millimeters and individually verified to ensure that no switching of identity had occurred. Fig. 2B shows a sample tracking in 2D; Fig. 2C shows tracked positions over time for the same recording as in 2B (see Suppl. Movie 1 for tracking results together with the original video). Behaviors of the mice were automatically scored for each frame by training multiple JAABA (Janelia Automatic Animal Behavior Annotator, (Kabra et al., 2013)) classifiers (Fig. 2B, line styles, see caption).

**Figure 2:**
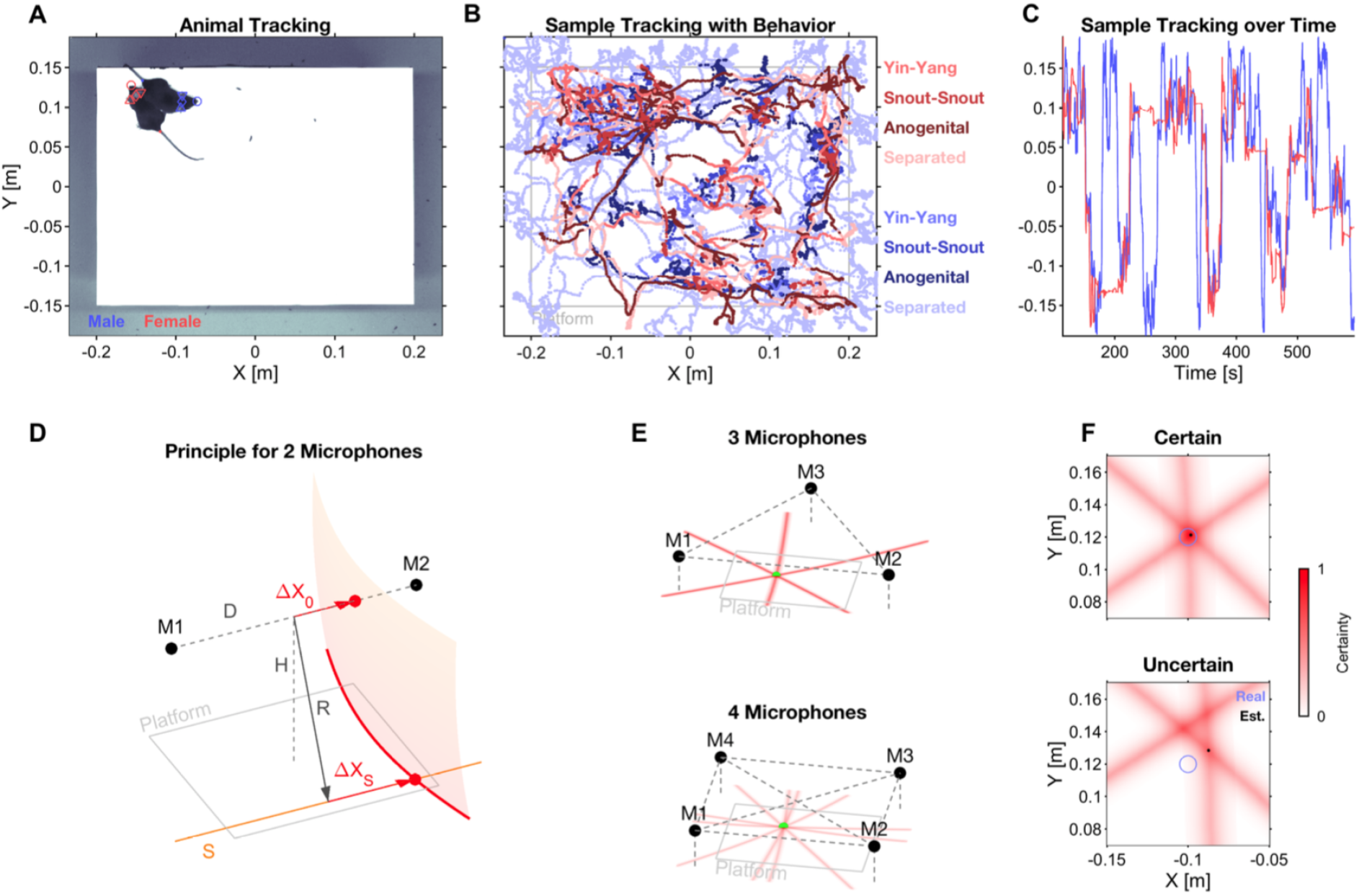
Localizing and attributing USVs to interaction partners in 2D. The assignment of USVs to a mouse requires (i, top row) visually tracking both mice and (ii, bottom row) localizing the origin of the USV in space, followed by assignment to the closest mouse (see *Methods* for detailed criteria). **A** Individual mice were tracked continuously at multiple body parts using DeepLabCut (Mathis et al., 2018), Male: blue, Female: red; Snout: circle, Left ear: downward triangle, Right ear: upward triangle, Head centre: rhombus, Tail onset: dot. Female mice were shaved on the top of their head to allow automatic recognition in DLC. **B** Sample trace from a single recording, including multiple automatically scored behaviors (JAABA, see (Kabra et al., 2013); indicated by thickness/brightness of the trace, see legend, color indicates sex). **C** Sample traces of snout markers of both animals over time after DLC tracking and post-hoc filtering, demonstrating smooth traces without switching. **D** Spatial localization of the origin of each USV was performed using multiple, partial localizations from pairs of microphones (M1/2), including correction for elevation. A cross-correlation measure (EWGCC; (Heckman et al., 2017)) indicated a set of possible origins between the microphones which lies on a 2D manifold (translucent red, right). Correcting for the height H of the microphones above the platform, a curve of possible *origin curves* (red, platform level) results for each single microphone pair (see *Methods* for details). **E** Complete localization in 2D can be performed using 3 or more microphones. The origin curves (light red) from each pair of microphones ideally intersect in a single point, indicating the unique origin of the USV (black dot). For 3 microphones (top, M1-3), there are 3 curves that can intersect. For 4 microphones (bottom, M1-4), there are 6 curves, which increases the accuracy and robustness against noise. **F** The certainty of localizing USV origin can be assessed on the level of single USVs. If the origin curves intersect in precisely one point, the certainty of the estimate is high, whereas the quality of the estimate is worse if various intersections of the curves are more spread out.

For the purpose of assigning USVs to their emitter, we developed a new USV localization technique, which improves the accuracy of spatial localization over previous techniques, achieving an MAE (median average error) of ∼12-14 mm (∼3-fold improvement in distance, ∼7-fold in area). The method takes our recently developed correction for microphone height (Heckman et al., 2017) and generalizes it to multiple microphones, improving accuracy (with the number of available microphones) and also allowing localizing sounds in 2D (using 3 or more microphones) or 3D (using 4 or more microphones). We refer to the generalized method as *SLIM* (Sound Localization via Intersecting Manifolds). Briefly, SLIM analytically estimates submanifolds (in 2D: surfaces) of a sound’s spatial origin for each pair of microphones (Fig. 2D) and combines these into a single estimate by intersecting the manifolds (lines, Fig. 2E). The intersection has an associated uncertainty which can be used to predict the precision of the localization estimate for individual USVs (Fig. 2F).

### SLIM substantially improves localization accuracy

We quantified the accuracy of SLIM for mice in social interaction (Fig. 3) both when the mice were in close proximity and far from each other. The position estimates aligned closely with the spatial position of the mouse’s snout that was closest to the estimate (determined from the video recording and the setup geometry, Exp. 1, Fig. 3A/B; for Exp. 2 with 3 microphones see Fig. 3 SD1). The one-dimensional accuracies in the left-right (MAE = 8.1 mm) and front-back direction (MAE = 8.4 mm) were comparable. Centered on the snout of the closest animal, the errors were distributed evenly in angle and decayed quickly with distance (Fig. 3C/D) with an MAE of 14.3 mm for all USVs (light green, see Fig. 3C/D) and 13.1 mm for the set of reliably assignable - referred to as ’selected’ - USVs (Fig. 3C/D, dark green, see Methods for details). The reliably assignable USVs constituted 84.3% of all USVs.

**Figure 3:**
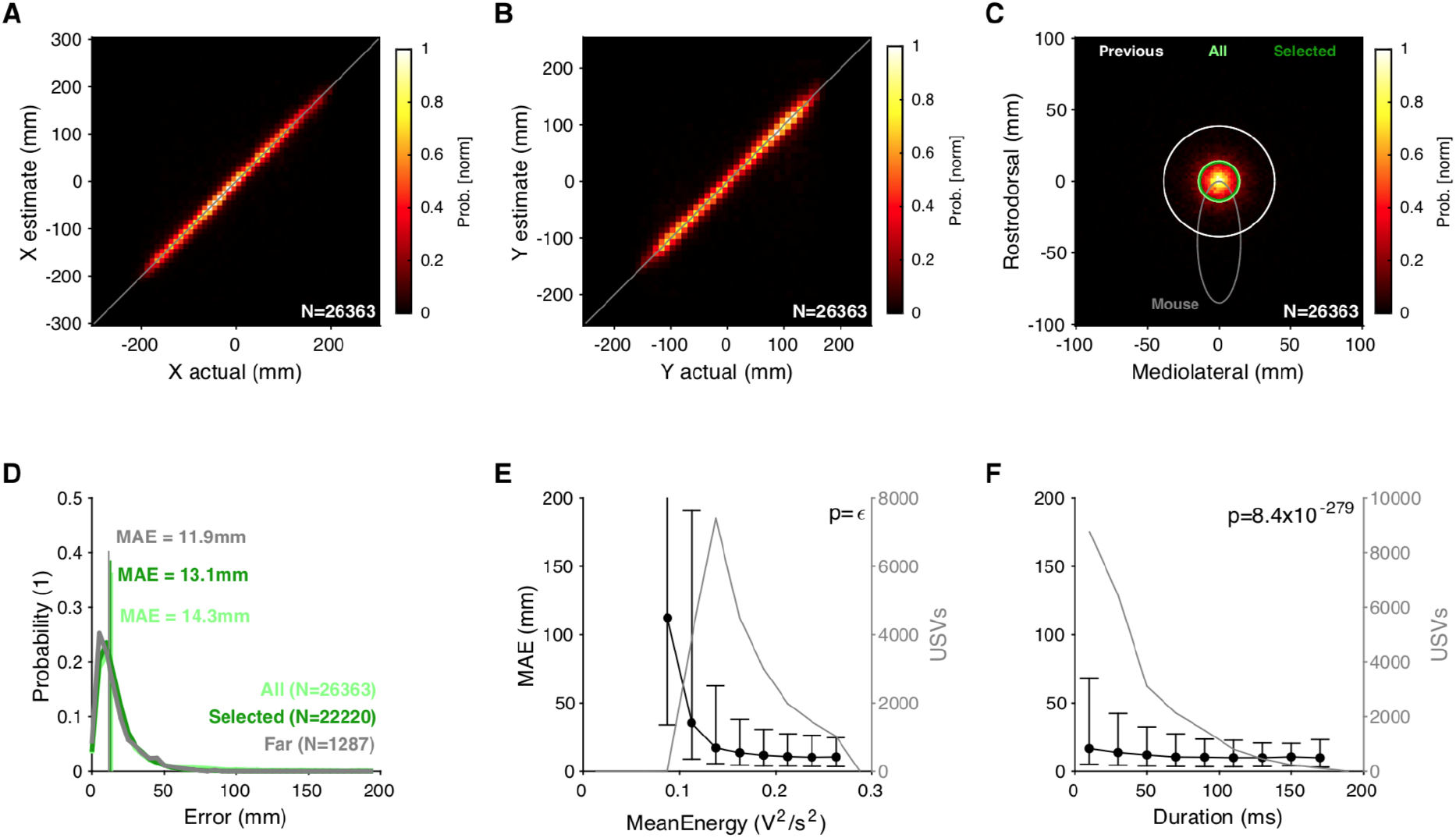
SLIM improves spatial accuracy in localizing vocalizations substantially over previous methods. **A** Density of actual (male) snout locations along the X dimension (horizontal in the video image) concentrated closely around the diagonal. Colours indicate peak-normalized occurrence rates. See Fig. 3 SF1 for accuracies obtained using a 3 microphone setup. **B** Same as A for the Y coordinate, again closely concentrated around the diagonal. **C** Combined 2D localization of USVs centered on the mouse snout. The average accuracy is visualized by a circle whose radius is the median average error (MAE; light green: all USVs; dark green: selected - i.e. reliably assignable - USVs see *Methods*; white: from (Neunuebel et al., 2015) for reference). **D** The distribution of localization errors for the entire set of vocalizations has an MAE of 14.3 mm (light green), while the reliably assignable ones (’selected’) had an improved MAE of 13.1 mm (dark green). When considering only USVs at times when the mice were >100mm apart, the MAE further improved to 11.9 mm. The error density is displayed as a normalized histogram. **E** Location accuracy significantly improved with the average acoustic energy of the vocalization (black). The detected vocalizations had a mean energy starting at 0.1 V^2^/s^2^ and ranging up to 0.3 V^2^/s^2^ (gray). The p-value was within the computer precision epsilon. **F** The MAE (black) showed a significant dependence on the duration of USVs, improving with longer durations, which are, however, more rare in comparison to short vocalizations (gray).

Assignment to the closest mouse can be erroneous as it is not based on ground truth data regarding which of the mice vocalized. To obtain a surrogate for ground truth data, we selected the USVs emitted when the mice were >10 cm apart from each other, i.e. much further than the estimated accuracy of the method. Here, the accuracy improved to an MAE of 11.9 mm (p=0.0002 for the selected, and p=3x10^-18^ for all USVs, Wilcoxon rank sum test) when only considering the reliably assignable USVs (Fig. 3D, gray). The improved accuracy for well-separated instances suggests that occlusion during close interaction may influence the localization estimate. Overall, these estimates compare favourably to the accuracy of previous methods also using 4 microphones (MAE = 38.6 mm, (Neunuebel et al., 2015), see *Discussion* for more detailed comparison). We also conducted a limited set of experiments with a single mouse on the platform which produced comparable spatial accuracy. As this condition was not suitable to reliably elicit a substantial number of vocalizations, it is not discussed in detail here.

As expected, the localization accuracy of SLIM was worse for low amplitude USVs (Fig. 3E, black, p<10^-100^, Kruskal-Wallis ANOVA) although those were infrequent in the overall set of USVs (gray curve, right axis). For USVs with high mean energies, the accuracy rapidly improved and stabilized at the highest energies at 10.9 mm. Furthermore, the localization accuracy showed a systematic dependence on duration by significantly decreasing from an average MAE of 17.1 mm for short USVs (0-20 ms) to an asymptotic accuracy of 10.2 mm for long USVs (>140 ms, p<10^-100^, Kruskal-Wallis test, Fig. 3F). In Fig. 3E/F, error bars indicate [5-95] percentiles instead of SEMs to display the variability rather than the certainty of the average.

In summary, SLIM provided reliable sound localization estimates with accuracies in the range of 10.2-17.1 mm depending on the USV’s intensity, duration, and relative animal position. Using the 4 microphone configuration, 84.3% of the USVs could be assigned and used for further analysis. The MAE for 3 microphones was ∼50% larger (see Fig. 3 SD1), which highlights the value of increasing the number of microphones (e.g. 8, as in (Sangiamo et al., 2020) to further improve the accuracy of SLIM).

### USVs are preferentially emitted in particular spatial relations which differ between sexes

In combination with automatic, deep learning-based dual animal tracking, SLIM allows us to investigate USV production during social interaction with high spatial precision. Specifically, we analyzed the relative spatial position of the animals in relation to USV density and spectral characteristics. The analysis in this section is based on all data from Exp. 1, i.e. Foxp2-R552H male, C57Bl/6J WT male and female combined. The next section separates the genetic variant from littermate controls on the male side.

For this analysis, the relative positions of the receiver animal are collected into an occurrence density map centered on the vocalizing animal’s snout direction, with the coordinate system appropriately translated and rotated for each USV. In this polar representation, the radial distance corresponds to snout-snout distance, and the angle describes the relative angle between the emitter’s snout direction and the receiver animal’s snout position (see illustration in Fig. 4A).

**Figure 4:**
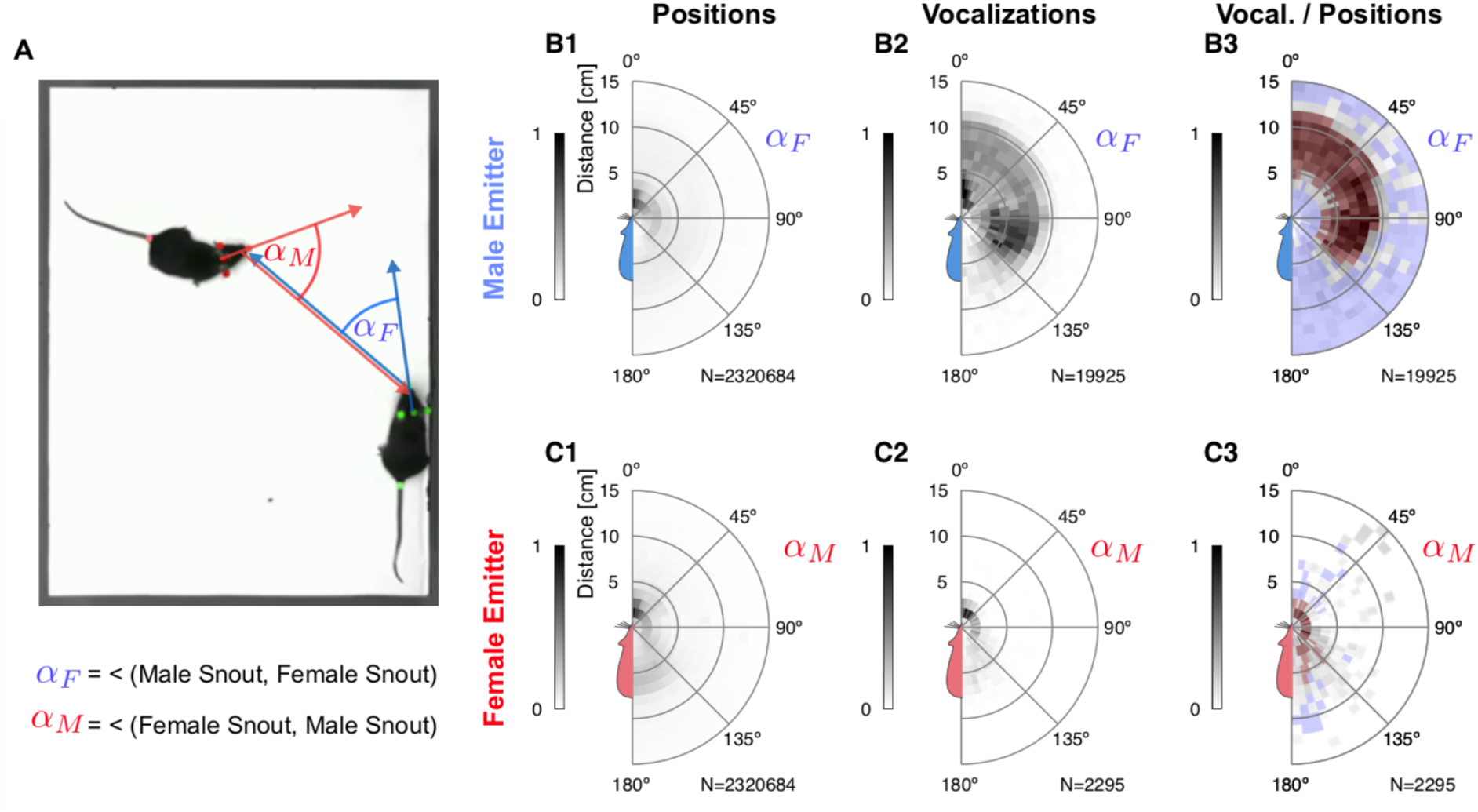
Vocalizations are emitted in particular spatial relationships and differ between sexes. **A** The relative position between the mice was analyzed in polar coordinates, with the distance between the snouts and the angle *α*_F/M_ between the head direction of the emitter and the head position of the receiver. Positive and negative angles (left and right side of the emitter mouse) are mirrored and averaged. **B1** First, the video data was used to establish the vocalization-independent spatial location prior distribution as an average across all video frames (N=2.3M). The female mouse was most frequently in facing, snout-snout contact with the male mouse (peak near 0° and 0-2 cm). **B2** USVs were emitted by the male mouse both in snout-snout contact but also very frequently when the female snout was about 5-10 cm away, which largely corresponds to snout-anogenital contacts. **B3** After normalizing by the relative position prior, (**B1**) USVs emitted in snout-anogenital location were the most abundant in the sense of a high conditional probability whereas snout-snout vocalizations in fact occur relatively infrequently. **C1** The male mouse was also predominantly in facing, snout-snout contact but also spent substantial amounts of time near the anogenital of the female mouse (light-maroon density between 140-180° and 5-8 cm). **C2** Female mice vocalized when the male was in snout-snout contact or behind/to the side of the female mouse. **C3** After normalizing by the relative position prior (**C1**), the most frequently encountered positions for females were the snout-snout, snout-anogenital (male at anogenital of female mouse), but also more distant relative positions. In **B** and **C** the colour displays peak-normalized histogram entries.

Mice predominantly vocalized when close to each other, i.e. within ∼10 cm of each other (Fig. 4B2/C2 and Fig. 1D). Overall, the vast majority of USVs was emitted by male mice (89.6%), however, female mice clearly vocalized as well (10.4%). After normalizing for their general relative position (Fig. 4B1/C1), we found that male mice vocalized most frequently when their snout was in close proximity to the female’s ano-genital region (Fig. 4B3, dark red arc within 5-10 cm, see also Fig. 4 SD1, showing the corresponding snout-to-anogenital densities). In contrast, female mice vocalized most when in snout-snout interaction, or when the male snout was close to the female’s ano-genital region (Fig. 4C3 and Fig. 4 SD1). Evidently, the relative spatial vocalization preferences of the animals differ substantially as their significant USV occurrence maps do not overlap (compare Fig. 4B/C3; p<0.05 for all bins, permutation test against spatially shuffled density values, red/blue hues indicate significant positive/negative deviation, respectively).

These salient differences in relative spatial position during vocalization between the sexes are likely mediated by behavioral contexts that present different motivational cues, e.g. tactile or olfactory.

### USV properties are shaped by relative position, genetics and sex

Exploiting the combination of high-accuracy localization of animals and vocalizations, we explored the influence of relative position, genetic variant and sex on the USV properties chosen by the mice (Fig. 5, and Fig. 5 SD1/2, respectively). Below, significances across groups (sexes/strains) are based on a 3-way, nested ANOVA analysis, with the predictors sex, genetic variant and individual animal, where the latter was nested w.r.t. to the first two.; significances across angles/distances and within group (sex/strain) on Kruskal-Wallis one-way ANOVAs; significances across group and angles/distances on regular 2-way ANOVAs (due to unavailability of a general, non-parametric 2-way test). All p-values and effect sizes (Cohen’s *D*) are reported in Fig. 5.

**Figure 5:**
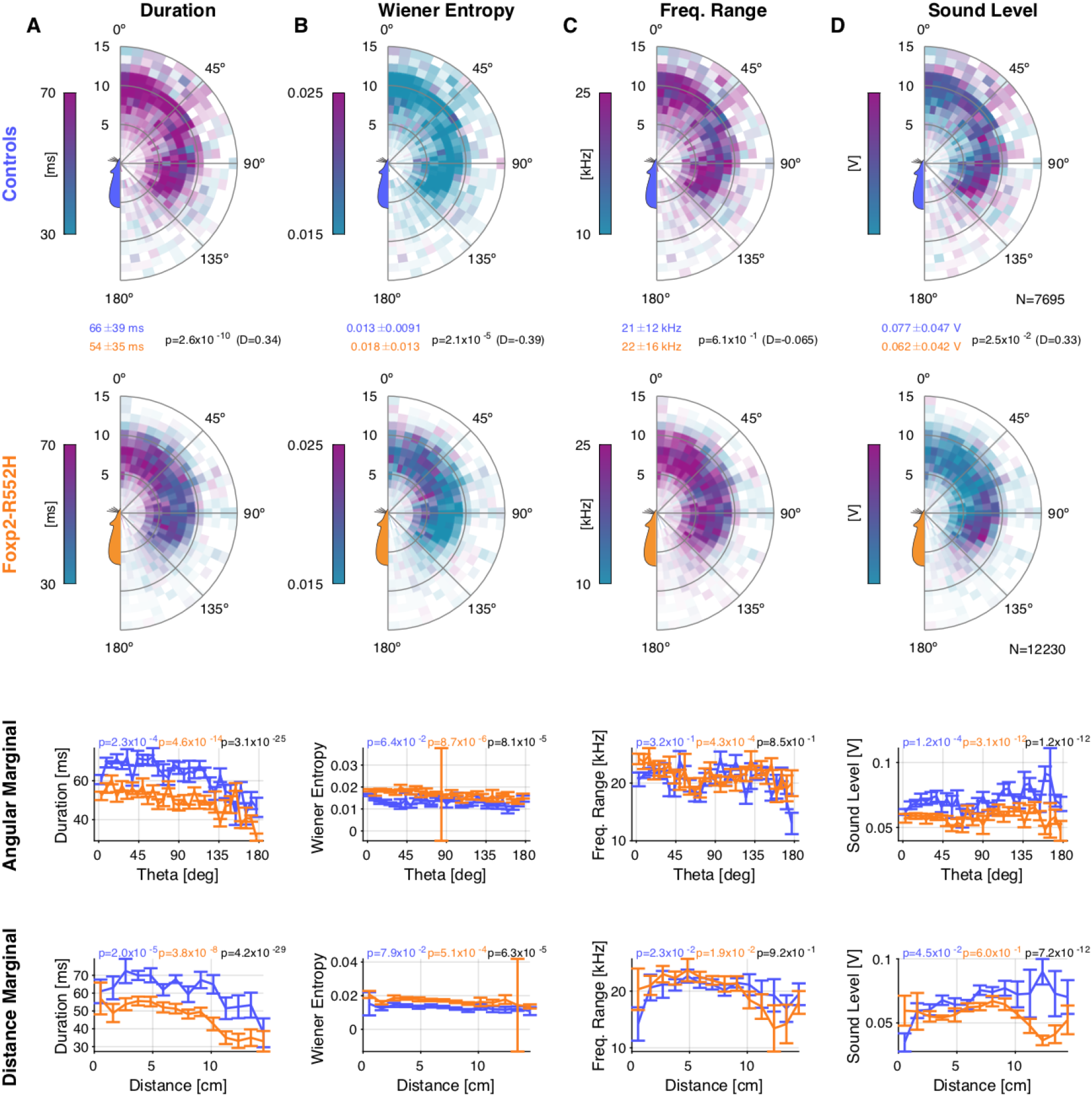
Foxp2-R552H mice produced USVs with different properties as a function of relative spatial location compared to their WT littermates. **A** The duration of male USVs was substantially longer for WT males than for Foxp2-R552H males (comparison of top two rows). Male USV duration depended on both angle and distance (bottom rows, controls: blue; Foxp2-R522H: orange). **B** The Wiener entropy of Foxp2-R552H USVs was greater than those of WT USVs, and dependent on both angle and distance. **C** The frequency range of Foxp2-R552H and WT did not differ strongly and showed mostly insignificant dependence on distance and angle. **D** The sound level of Foxp2-R552H USVs was smaller than that of WT USVs, in particular for distances >10 cm and angles >100°, indicating snout-anogenital contact with the animals facing in opposite directions. In the top two plots, hue indicates the respective average USV property in a given location while the opacity indicates the density of USVs (using the spatial vocalization density; see Fig. 4, last column). The hues do not correspond to the colors of the strains. Significances for all comparisons are given in the figure, i.e. Wilcoxon tests between groups (between first two rows, in addition to mean ± s.d. for each group); for the bottom two rows Kruskal-wallis ANOVA tests for within group and across angle/distance (coloured according to group); 2-way ANOVA significance for combined group and angle/distance comparison (black).

We collected the average properties of USVs emitted by a given group in relation to the interacting group at the time of vocalization in a combined colour-density plot (Fig. 5, top two rows). In these plots, the intensity indicates the density of occurrence of USVs in the relative spatial bin while the colour hue indicates the property value, e.g. a USV’s duration. More intense colours thus also correspond to more reliably estimated means in this location.

The duration of USVs emitted by Foxp2-R552H was significantly shorter (Fig. 5A, Foxp2-R552H (orange): 66 ms, WT (green): 54 ms, p=2.6x10^-10^). In addition, the duration of USVs decreased with angle from front to back (p<10^-3^ for both) and with snout-snout distance for both groups (p<10^-3^ for both) and differed significantly also on the angle and radius marginal (p<10^-24^ for both). As above, these interactions are largely snout-anogenital (directly verified in Fig. 4 SD1).

The *Wiener entropy* (Ivanenko et al., 2020; Johnston, 1988) of USVs also differed significantly (Fig. 5B, Foxp2-R552H: 0.018, WT: 0.013, p=2.1x10^-5^), with Foxp2-R552H mice emitting USVs that exhibited higher Wiener entropy, in particular when the male was behind the female (p=8.7x10^-6^), i.e. in snout-anogenital but also during snout-snout interactions. Since the spatial location of elevated Wiener entropy in female mice is a combination of radial and angular ranges, the marginal distributions only show a reduced effect. Further, the Wiener entropy in Foxp2-R552H and WT mice differed in their dependence on angle (p=8.1x10^-5^) and radius (p=6.3x10^-5^).

The frequency range of the USVs did not show a significant difference between Foxp2-R552H and WT mice overall (Fig. 5C, Foxp2-R552H: 22 kHz, WT: 21 kHz, p=0.61) and also behaved quite similarly for the angle or distance for either genetic variant.

The levels at which USVs were produced differed significantly between Foxp2-R552H and WT mice (Fig. 5D, Foxp2-R552H: 0.062V and WT mice: 0.077 V, p=0.025; given in s.d. of microphone output voltage because translation to local level in dB is highly uncertain), with WT mice vocalizing at substantially higher intensity compared to Foxp2-R552H mice. Both the sound level of Foxp2-R552H and WT USVs showed a significant dependence on angle, while the distance did not significantly influence the sound level (see Fig. 5D for specific p-values).

We also conducted an analogous analysis comparing WT males and females. This analysis also indicated significant differences between sexes (see Fig. 5 SD1), although the small number of female USVs leads to rather sparse densities. Overall, male C57Bl/6 WT USVs were significantly longer, had a larger frequency range and a lower Wiener Entropy than female C57Bl/6 WT mice. In summary, in dyadic male-female interactions, the relative spatial position, genetic variant and sex all had a significant influence on various properties of USVs chosen by the mice.

### Relations between spectrotemporal properties, sex, strain, and relative position

Lastly, we aimed to disentangle the relation between the spectrotemporal properties of USVs in relation to their emitter’s sex, strain, and relative spatial position. For this purpose, we applied UMAP dimensionality reduction (McInnes et al., 2018) to a set of USV properties (Fig. 6A, see *Methods* for details). Projected to 3D, the spectrograms grouped into an intricate spatial arrangement with structure on both the macro and micro level (Fig. 6C). Post-hoc classification into spatial clusters (k-means) indicated that on the order of 100 clusters would be needed to account for the substructure although many of the clusters were not clearly separated (see Suppl. Movie 2 for a rotating version of this plot). These results indicate that previous classification schemes into a handful of clusters may need to be revised.

**Figure 6:**
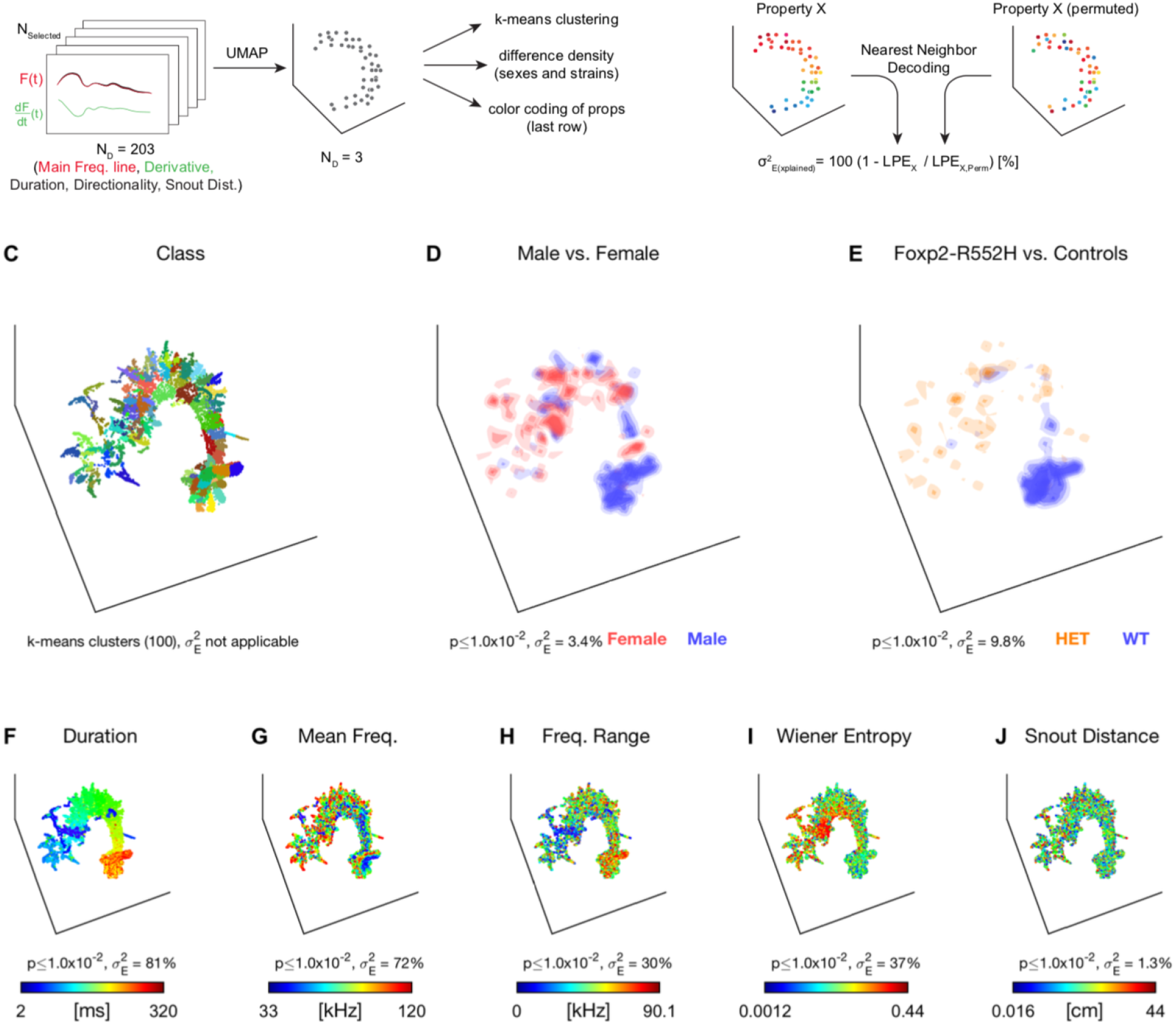
Dimensionality reduction of USV properties detects relations between phonetics, strain, sex, and relative position. Supplemental Movie 2 shows the same data revolving in 3D, resolving depth ambiguities. **A** Each USV was represented by its main frequency line (up to 100 ms), its derivative, the duration, the directionality in frequency, and the snout-to-snout distance (203 dimensions) and then reduced to 3 dimensions using UMAP (McInnes et al., 2018). The analysis was based on the set of selected vocalizations (see Fig. 4). The results are differently colour-coded for each property. **B** We evaluated whether a property of the USV was related to others after dimensionality reduction by comparing the original (left) to shuffled data (right; same spatial and value distribution) using nearest neighbor decoding. This allowed us to assess the percentage of variance explained as the relation between the local prediction error (LPE, see *Methods* for details) of the original and shuffled data. **C** After dimensionality reduction, the set of spectrograms exhibited a rich structure. We highlight the richness of the substructure here by running clustering (k-means, k=100, different colors). However, clusters are often not clearly separated but rather connected. The properties in the following panels partially clarify the origin of this substructure. **D** Sex showed a significant contribution to accounting for the neighborhood structure, shown here via local density differences between male and female emitters. While significant, these differences explain only 3.4% of the LPE. **E** The Foxp2-R552H variant (male vocalizers only) also significantly accounted for the local structure, accounting for 9.8% of the LPE. Interestingly, differences in spatial density between the strains (Foxp2-R552H: orange; WT littermate: green) partly coincided with the sex differences in **D**. **F** Among the tested properties, duration best explained the neighborhood structure of the spectrograms by being able to account for 81% of the LPE. **G** Mean frequency also explained a substantial part of the neighborhood structure, although largely locally ’orthogonal’ to the duration (i.e. multiple local gradients of frequency), accounting for 72% of the LPE. **H** Frequency range explained 30% of the LPE and appears correlated in spatial distribution with duration. **I** The Wiener Entropy also explained a substantial part of the LPE (37%). **J** Different snout-snout distances only contributed borderline significantly to the structuring accounting for 1.3% of the LPE. Supporting figures 6 SD1/2 and Supplemental Movies 3/4 show the same analysis when excluding duration or duration and mean frequency.

We further analyzed the data’s neighborhood structure by associating it with a range of properties associated with each spectrogram, e.g. its emitter’s sex, Foxp2-R552H genetic variant, relative position, and spectrotemporal properties. The degree to which a given property contributed to explaining the spatial structure was analyzed using nearest neighbor prediction on original and permuted datasets (Fig. 6B, see *Methods*). This analysis yielded a measure of explained variance 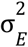 defined as one minus the fraction of Local Prediction Errors (LPE, see *Methods*) for the original dataset and the average of multiple permutations. Sex and Foxp2-R552H (male only, Foxp2-R552H vs. controls) significantly contributed to explaining the neighborhood structure of USVs, accounting for 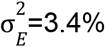 (p<0.01, Fig. 6D) and 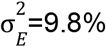 (p<0.01, Fig. 6E), respectively. Sex exhibited spatially localized differences in the contribution of female and male vocalizations (shown as red and blue spatial regions, respectively, in Fig. 6D). Similarly, the genetic variant exhibited differences in the spatial density of vocalizations (shown as orange (Foxp2-R552H) and green (controls) spatial regions, respectively, in Fig. 6E).

The largest contributions to explaining the spatial structure were made by spectrotemporal properties, i.e. a USV’s duration, mean frequency, frequency range, and (mean) Wiener entropy. The duration explained 82% of the LPE (Fig. 6F). The *Mean Frequency* (Fig. 6G), *Frequency Range* (Fig. 6H), and *Wiener Entropy* (Fig. 6I) individually explain 72%, 30%, and 37% of the LPE, respectively. Note, that these contributions do not need to sum to 100% as they can be correlated with each other and explain the same structural similarities between USVs. The snout-snout distance also explained a significant part of the structure (1.3% LPE, p<0.01, Fig. 6J).

The structural similarity analysis indicates that while USV spectrograms are predominantly grouped by their spectrotemporal similarity (Duration, Mean Frequency, Frequency Range, Wiener Entropy), sex- and in particular genetic variant-differences can explain part of the spatial structure, indicating sex- and Foxp2-R552H-specific differences in vocalization properties. Similarly, snout distance made a significant contribution though less than strain and sex.

## Discussion

In the present study, we combined a novel acoustic spatial localization method with state-of-the-art animal tracking to obtain a higher level of accuracy in localizing and assigning sounds to their emitter. The resulting spatial maps indicate that vocalizations differ depending on relative spatial location, sex, and genetics. The present method generalizes to 3D localization and a larger number of microphones and thus provides a versatile tool to study other strains and/or species in close social interaction, such as avian species.

### Differences in vocalization between male and female mice during social interaction

Previous studies have investigated the relation between vocalization properties and interaction types, albeit at lower spatial accuracy. While an earlier study found no significant dependence of a subset of properties (duration, interval) on behavior (Hammerschmidt et al., 2012), we recover such a dependence in the near-field interactions between the mice. Specifically, we find the relative position and the sex to influence the USVs chosen by the mice, with respect to their duration, Wiener entropy, and frequency range.

There have been earlier studies (Heckman et al., 2017; Warren et al., 2018) which have found sex-dependent differences in vocalization properties. Some findings that appear conflicting may be attributable to different strains and experimental setups in the studies, both of which are factors that may influence vocalization behavior (Heckman et al., 2016). Considering the dependency of vocalization properties on the spatial relation between the interacting mice shown in the present study, future research may include this factor as well when comparing USV properties between the sexes.

Our observation that vocalization likelihood is linked to relative body position of the two interacting mice (Fig. 4B3/C3) is consistent with the findings of Neunuebel et al. (Neunuebel et al., 2015). To investigate any potential causal relationship between vocalizations and behavioral changes, further studies with greater focus on the interpretation of behavioral states would be needed.

A recent study (Sangiamo et al., 2020) demonstrated an influence of vocal expression on the behavior of interacting animals in a way that is consistent with the present findings although the spatial relations analyzed here and behavioral interaction types analyzed there cannot simply be mapped onto each other. We did not perform a classification into vocalization types as the focus of our study was on improving spatial localizations and because of the potential overlap between the extracted categories.

While females can vocalize as frequently as male mice in other social contexts, e.g. resident-intruder interactions (Hammerschmidt et al., 2012; Ivanenko et al., 2020), male mice are the main vocalizers in social interactions. The exact fraction of USVs emitted by females as concluded in all previous studies on dyadic courtship has varied, ranging from 18% (Neunuebel et al., 2015), 17.5% (Sangiamo et al., 2020), and 16% (Heckman et al., 2017) to 10.5% in the present study (N.B., (Warren et al., 2018) and (Warren et al., 2020) did not indicate fractions of female vocalizations to our knowledge). This variability is likely attributable, in part, to differences in the precise paradigm (duration, number of animals, environment, etc.), strains, and individual mice. However, the precision in localizing USVs is very likely another contributor. As the present data suggests, female mice are more likely to vocalize during close snout-snout interactions (Fig. 4C3). Imprecise localization will therefore affect the attribution to female animals more strongly than to males, whose attributions during snout-ano-genital interactions remain largely unaffected (Fig. 4B3). One consequence of imprecise localization would in that case be that male vocalizations are erroneously assigned to the female, which would bias their fraction upwards. In future studies, a higher spatial precision in localization should help disentangle the cause of these varying female vocalization rates.

### Differences in vocalization between Foxp2 mutants and wildtype mice during social interaction

Rare heterozygous mutations that disrupt the human *FOXP2* gene have been implicated in a developmental speech and language disorder, leading to studies of functions of its orthologues in a range of other species (Fisher and Scharff, 2009). Disruptions of mouse *Foxp2* have been linked to changes in murine vocalization behavior in several reports (Castellucci et al., 2016; Gaub et al., 2010; Shu et al., 2005). Most of the early studies that point out this linkage focused exclusively on mouse pups, and there were inconsistencies noted between different reports (French and Fisher, 2014; Groszer et al., 2008). Although USV sequence length has previously been shown to be affected in adult mice with heterozygous *Foxp2* disruptions (Castellucci et al., 2016; Chabout et al., 2016), evidence for mutation-related changes in the USV sound structure or syllable repertoire remains inconclusive. Various studies compared vocalizations of adult mice with a heterozygous *Foxp2* mutation to their wildtype littermates and found there to be no detectable difference in acoustic parameters like USV duration, mean frequency or amplitude (Castellucci et al., 2016; Chabout et al., 2016; Hammerschmidt et al., 2015). It has to be noted, however, that these studies did not take into account the spatial context in which those USVs have been produced.

Gaub *et al*. (Gaub et al., 2016) considered the spatial relations implicitly by analyzing vocalizations grouped by interaction types, e.g. mutual sniffing, genital sniffing, head sniffing, and others. For example, for USV duration, their study finds significant differences between interaction types, but not between WT and heterozygous Foxp2-R552H mice. In our study, we resolve the interactions spatially instead and find significant differences in a number of properties. Here we demonstrate that male Foxp2-R552H mice exhibited shorter and quieter USVs, with higher Wiener entropy as compared to male WT littermates, but overall showed similar dependence on angle and distance. Additional differences probably exist for other properties. However, an exhaustive treatment of these properties was not the focus of the present study. We hypothesize that this difference in results to the study by Gaub *et al*. (2016) is due to the improved spatial resolution in our analysis, which, in addition to spatially resolved interactions, allows a more reliable assignment of USVs to their emitter.

### Methodological advances over and comparison with previous studies

The precision of spatial localization is key for studying the complete communication of mice during social interaction, as a high accuracy enables a reliable assignment even in close interaction. The technical innovations -explicit correction for microphone height, envelope weighted cross-correlation, and the combination of an arbitrary set of microphone pair estimates to a single estimate- introduced by SLIM provide a high accuracy for spatially localizing vocalizations. To our knowledge, this accuracy (11.9-14.3 mm) is substantially higher than in previous studies (i.e. compare Neunuebel et al. 2015: MAE = 38.6mm). However, some recent studies (Sangiamo et al., 2020; Warren et al., 2018) using 8 microphones have focused on the percentage of assigned USVs and not reported average errors in distance of assignment for either single or multi-animal tracking. While we therefore cannot quantitatively compare the spatial accuracy directly, we can partially compare the percentage of USVs assigned. In our study, 84.3% of USVs were reliably assigned, compared to 40.4% in (Neunuebel et al., 2015), 64% in (Sangiamo et al., 2020), and 76% (4 Mics) / 92% (8 Mics) in Warren et al. (2018, Table 1), with the latter using a Jackknife estimate. The results with 8 microphones from Warren et al. (2018) are impressive and demonstrate the advantage of making use of a larger number of microphones, and we expect that both accuracy and assigned fraction in SLIM would further improve under such circumstances. Precise quantitative comparison between these studies and ours is, however, complicated by the fact that their study was performed with 4 mice at a time, instead of the 2 used in the present study. While this will make it harder to pass the mouse probability index MPI>0.95, there are more opportunities/animals to which a localized USV can be assigned, making it hard to compare the fractions. Further differences include the acoustic properties, the size of the interaction space, and the techniques for extracting USVs.

### Limitations of the current approach

A particular challenge for assigning USVs during social interactions are snout-snout interactions where the potential acoustic sources are closest to each other, roughly within 20 mm. Our results suggest that it is particularly this type of interaction where female mice choose to vocalize most frequently (see Fig. 4). We ran a simulation (Fig. 3, SD3) which indicates that after the MPI selection procedure, the accuracy of our approach stays very high (∼84.3%) even for this closest interaction, thus rendering the interpretation of the close interaction results trustworthy.

While the automated tracking was largely quite accurate, residual tracking errors contributed to the estimated precision of SLIM. We hand-tracked a small subset of recordings and noted a further improvement in localization accuracy on the order of 1-2 mm. For the present study, we chose not to hand-track all recordings as this would have been unfeasible for all frames (>2M) needed for computing the conditional spatial densities of vocalization. Subsequent improvements in animal tracking will be required to further reduce the acoustic tracking and assignment errors.

Lastly, a challenge of laboratory conditions is the unnatural setting in which the animals are brought together, which includes a relatively small area, acoustically insulated walls, and a short time of acclimation for the animals to this new environment (partly to prevent mating during the experiment). These factors may influence the anxiety level of the animals and thus also their vocalization behavior, e.g. vocalization rates, choice of syllables, and sequencing. Transitioning to a more natural environment would be beneficial, but on the other hand additional objects in the space (e.g. shelters, rustling of nesting material) could negatively affect the precision of sound localization. In addition, it would be ideal to transition to a more continuous monitoring of mice in order to study them under conditions that are less stressful, which would likely increase vocalization rates (similar to Neunuebel et al. 2015).

### Conclusions and future improvements

Precise spatial localization of vocalizations and thus reliable attribution to their emitter opens new possibilities for studying social behavior, such as automatically monitoring the well-being of animals in laboratory cages and designing new closed-loop feedback paradigms. Further improving the accuracy to master the most challenging situations with sound sources in extremely close proximity, e.g. in snout-snout contact, will require more microphones and further refined analysis techniques, for example those that combine visual (pose) and acoustic information, e.g. a deep neural network that processes video and audio streams in parallel with the goal of learning to take occlusion into account. A fruitful research direction here is virtual acoustics as a basis for creating large-scale datasets for deep learning (Menzer et al., 2011).

## Acknowledgements

The authors would like to thank Ron Engels for help in constructing the setup and Sebastian Tiesmeyer for helping in setting up DLC. This work was supported in part by the European Commission (Horizon2020, nr. 660328). SEF and CJLMS were supported by the Max Planck Society. Funding for equipment and animal costs was provided by a Technology Hotel Grant (ZonMW, 40-43500-98-4141). BE was supported by an NWO-VIDI Grant (016.VIDI.189.052).

## Materials & Methods

All experimental procedures were approved by the animal welfare body of the Radboud University under the protocols DEC-2014-164 and DEC-2017-0041 and conducted according to the Guidelines of the National Institutes of Health.

### Animals

Two distinct groups of mice were used in this study (see Table 1 for overview). The first group consisted of 12 Foxp2-R552H males as well as 12 male and 4 female wild type littermates on a C57Bl/6J background (Groszer et al., 2008), referred to as C57Bl/6J WT, or ’controls’ for the male mice. The second group consisted of 10 male and 2 female CBA/CaJ WT mice. The animals were 8 weeks old at the start of the experiments. After 1 week of acclimatization in the animal facility, the experiments were started. Mice of the same sex and strain were housed socially (2-5 animals per cage) on a 12h light/dark cycle with ad libitum food and water in individually ventilated cages.

### Recording Setup

The behavioral setup consisted of an elevated platform inside a sound-insulated booth, together with multiple ultrasonic microphones and a high-speed camera.

The booth had internal dimensions of 70x130x120 cm (LxWxH). The internal side walls and the floor of the booth were covered with acoustic foam (Thickness: 5 cm, black surface Basotect Plan50, BASF), which – according to the product specifications – shields against external noises above ∼1 kHz (sound absorption coefficient > 0.95 defined as ratio between absorbed and incident sound intensity; corresponding to >26dB shielding in addition to the shielding provided by the booth). Additionally, the foam eliminates internal reflections of high-frequency sounds, in particular USVs. Illumination was provided via three dimmable LED strips mounted to the ceiling, providing light from multiple angles to reduce shadows.

The interaction platform was constructed from slotted aluminum (30x30 mm) covered by a 40x30 cm rectangle of acoustic foam (thickness 5 cm, Basotect Plan50, white surface, with a laminated surface to simplify cleaning feces), with its surface located 25 cm above the floor (i.e. 20 cm above the foam on the booth floor). The platform had no walls to avoid acoustic reflections and was located centrally in the booth.

Sounds inside the booth were recorded with three or four ultrasonic microphones (CM16/CMPA48AAF-5V, flat (±5 dB) frequency response within 7-150 kHz, AviSoft, Berlin) at a sampling rate of 250 kHz. An analog low-pass filter at 120 kHz prevented aliasing and excluded contributions beyond 120 kHz. Recorded data was digitized using a data acquisition card (PCIe-6351, National Instruments). The microphones were placed in well-defined locations around the platform (see Fig. 2 for visualization). In the 3 microphone setup, the placement was in a triangle which contained the platform. In the 4 microphone setup, the placement was in a rectangle that contained the platform. In both cases, the microphones were placed at a height exceeding the platform (+13.3 cm for 3 microphones and +12.1 cm for 4 microphones) to reduce sound being blocked by the animals during interaction. The position of the microphone was defined as the centre of the recording membrane. The rotation of the microphones was chosen such that they aimed at the centre of the platform. To maximize the captured sound based on the microphones’ directional receptivity (∼25 dB attenuation at 45°), the microphones were placed away from the corners of the platform, i.e. 5 cm in the long direction (40 cm) and 6 cm in the short direction (30 cm) of the platform.

The camera was mounted centrally above the platform at a height of 123.5 cm (measured from the front end of the lens) w.r.t. the floor of the box, i.e. 98.5 cm above the platform surface. Video was recorded with a field of view of 46.9 x 37.5 cm (Lens: 12.5 mm, Cosmicar) at ∼50 fps and digitized at 640x512 pixels (effective resolution of ∼0.733 mm/pixel; Camera: PointGrey Flea3 FL3-U3-13Y3M-C, Monochrome, USB3.0). The shutter time was set to 10 ms to optimize illumination.

The interaction platform, the camera, and the microphones were mounted to a common frame made from slotted aluminium to guarantee precise relative positioning throughout the experiment. The frame was mounted to the floor of the booth.

### Experimental Procedures

Each experiment consisted of free interaction between a male and a female mouse on the platform. The female mouse was placed on the platform first and, shortly after, the male mouse was added. The recording was started before placing the female mouse and continued for 8 min. The recording was only interrupted if one of the mice jumped from the platform, which occurred in <5% of the recordings. The sequence of the animals was randomized daily.

Overall, we performed two very similar sets of experiments, denoted as follows:

1. Social interaction of a male C57Bl6/J WT or Foxp2-R552H mouse with a C57Bl6/J WT WT female mouse using 4 microphones (females identifiable by shaved spot on the head).
2. Social interaction between a pair of unmarked CBA/CaJ WT mice using 3 microphones (females identified by first arrival on platform).

the results, we separate these experiments where appropriate, e.g. to quantify the difference in accuracy for 3 or 4 microphones or vocalization/behavioral differences across strains. As only the female in set #1 was marked, we only use automatic, all-frame tracking for this set (see below).

### Data Analysis

The data analysis involved multiple stages, including animal tracking, detection, automatic localization of USVs, and behavioral analysis, all described in detail below. The code for the data analysis is made available together with this publication in an open repository upon acceptance (https://data.donders.ru.nl/collections/di/dcn/DSC_626840_0006_717).

### Animal Tracking from Videos

For the recordings from Exp. 1, mice were tracked offline in the XY-plane using DeepLabCut (Mathis et al., 2018) using multiple markers. For the recordings of Exp. 2, manual tracking of the snout and head centre of both animals was performed at the temporal midpoint of each vocalization.

For automatic tracking using DLC, a training set was created (1200 frames) containing manually placed markers for five locations for each animal, i.e. snout, head centre, ears, and tail-start. DLC was then trained with this data (DLC v.2.1, running on a GTX 1070 GPU with NVIDIA driver version 390.77, on Ubuntu 18.04.1 LTS). The resulting neural classification network was then used to predict marker locations for all frames in all recordings. Visual inspection revealed that the results were generally quite accurate for Exp. 1, where the female mouse had been labeled by shaving a small region on the head. Occasional jumps in markers were corrected with the use of a post-processing script, which used a custom set of median filters and short-range interpolation. Subsequently, we obtained clean trajectories of both animals (see Fig. 2B/C for a sample tracking).

For 9 recordings from Exp. 1, shaving a spot on the head was insufficient to provide good separation between the animals. We used an alternative strategy to track animals in these recordings: We requested in DLC multiple estimates (10 candidate locations for each feature) and performed custom linkage of body parts of the same animal between subsequent frames. Briefly, the strategy was as follows: Candidate locations for each body part were clustered (k-means), averaged, and then analyzed spatially to determine whether they could belong to the same mouse. Cluster averages that were safely attributed to a single mouse were taken to be the body part location. This was typically the case when the mice were spatially separated. Frames in which this was successful for at least one body part were then used as starting points for linking closeby candidate locations of neighboring frames in an iterative fashion (for details, see `C_trackMiceDLC.m` and tracking data provided in repository). Successful identification was confirmed by visual inspection for all recordings.

Manual tracking of recordings in Exp. 2 was performed by multiple human observers (GOS, JH, DL, BvR, AvdS). They were presented with a combined display of the vocalization spectrogram and the concurrent video image for the temporal midpoint of each vocalization (custom-written, MATLAB-based visualization tool). Users could freely scroll in time to identify female and male animals. For time efficiency, only the snout and the head centre (mid-point between the ears) were manually tracked. These points define a vector indicating the head location and gaze direction, which was required in subsequent behavioral analysis.

### Behavioral Analysis

To classify animal behavior, a machine-learning-based annotation system was used (Kabra et al., 2013). Based on visual observation of the most distinctive behaviors, we trained three classifiers, each annotating a single type of behavior, namely (1) close snout-snout interaction, (2) close male-female chases, and (3) mutual snout-at-tailbase (’Yin-Yang’) body contact. The classifier for the first behavior class was trained on 2,968 manually labeled frames (982 positive examples and 1,986 negative examples), the second on 1,644 (648 positive examples and 996 negative examples), and the third on 1,827 (609 positive examples and 1,218 negative examples). The accuracy of automatic annotation was evaluated on a set of manually labeled frames excluded from training. For the case of snout-snout interaction, this set contained 4,903 frames; for male-female chases, 11,872 frames; for snout-at-tailbase body contact, 4,513 frames. The three classifiers had false-positive and false-negative rates of respectively, < 0.1 % and 11.3%; 0.4% and 8.9%; 4.6% and 8.0%. Sample data from the JAABA classifiers is shown in Figure 2B (different line-styles indicate different behaviors, see figure caption).

#### Detection of Ultrasonic Vocalizations

Mouse USVs were detected automatically using a set of custom algorithms (see `VocCollector.m`) described previously (Ivanenko et al., 2020). A vocalization only had to be detected on one microphone to be included into the set. In total, we collected 26,363 USVs from 93 recordings in Exp.1 and 11,729 USVs from 79 recordings in Exp. 2.

#### Localization of Ultrasonic Vocalizations

The spatial arrangement of the current microphones allows spatial localization of sounds in two dimensions. Temporal differences between the microphones provide the most precise estimate (∼1.37 mm, for 4 µs = 1 sample at 250 kHz, based on the speed of sound in air).

We here introduce a localization technique for three dimensions, generalizing the analytical approach introduced in (Heckman et al., 2017). Briefly, for each pair of microphones, a curved surface of candidate locations is computed. These surfaces are then intersected with each other and the ’snout plane’ to obtain a density of candidate locations in this plane of social interaction. Finally, a single point estimate is formed from this density, including its associated spread as a measure of confidence for each vocalization estimate (see below for details).

First, we employed envelope-weighted generalised cross-correlation (EWGCC, for each pair of microphones: 6 pairs in Exp. 1, and 3 pairs in Exp. 2; (Heckman et al., 2017)). We extracted the peak of each EWGCC to estimate the most likely time delay Δ*T* for every vocalization. If the vocalization emanated from the line connecting the microphones, the location could be easily computed as the distance from the midpoint between the microphones Δ*X*_0_ = *v_sound_* Δ*T*/2(see Fig. 2D, red arrow). However, generally, the vocalization will not emanate from the line connecting the microphones.

We can compute all candidate locations in 3D space surrounding the microphones (see Fig. 2A for illustration), via (for derivation see (Heckman et al., 2017)

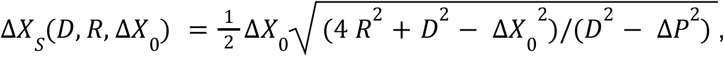

where *R* (Fig. 2D, dark gray, see below) is the distance between the line *L* connecting the microphones (Fig. 2D, gray dashed) and a given 1D subspace*S*(Fig. 2D orange) parallel to *L*, and*D* is the distance between the two microphones.Δ*X_S_* (Fig. 2D, red vector) is then the distance inside*S* from the plane orthogonal to*L*located at the midpoint between the microphones. Iterating this for all subspaces S provides a 2D surface of candidate locations (Fig. 2D, shaded surface) defined by the following relation:

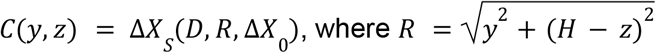

where *y* and *z* are measured from the centre of the platform, i.e. *C* gives the *x* coordinate for a combination of *y* and *z* (defining the above mentioned subspace *S*). In the interaction plane, this surface intersects as a curved line, referred to as an *origin curve* (Fig. 2D, red).

As this is still relative to the coordinate system aligned with the two microphones, the surface has to be appropriately rotated in the *xy*-plane to the actual two microphone positions. This is performed by a basic rotation matrix, with the angle defined by the angle of the connecting line between the microphones and the default coordinate system.

For each USV the above localization was performed on multiple, overlapping subwindows (length: 60 ms, moved in steps of 3 ms). The final localization was then computed as the median of the localization separately across dimensions for all sub-window estimates with an estimated localization accuracy (see below) of less than 40 mm (taking inot account the scaling below).

Lastly, estimates that fell outside of 50 mm from the platform edge on either side were projected onto this surrounding rectangle (platform edge plus 50 mm), as it was known that USVs could not originate from further out. In total, ∼0.5% of estimates were corrected in this way.

*Localization Accuracy* The quality of single localization estimates varies with each vocalization’s signal-to-noise ratio, frequency content, and representation across the four microphones. Knowing the quality per vocalization is a useful selection criterion, in particular if high precision is required during close interactions. We define the location accuracy *LA* as the standard deviation of all locations, >90% of the maximum of the intersection density of all origin curves (see e.g. Fig. 2F). *LA* is correlated to the accuracy of a given localization, but *LA* does not describe the standard deviation of the localization errors per se. Hence, it allows for a scaling factor to relate it to the actual errors, set to 4 in the present analysis.

### Spatial Vocalization Analysis

The acoustically estimated origin of a vocalization was then related to the candidate locations of the two mice in the corresponding frame obtained from the visual tracking. For each mouse, we used a position on the line connecting the snout to the head centre as the most likely origin of vocalization. The best overall match between acoustically and visually estimated positions was obtained at a distance of 10% from the snout for handtracked recordings, and a distance of 40% for automatically tracked recordings, probably reflecting differences in the detailed marker locations between the tracking strategies. Following the approach of (Neunuebel et al., 2015), we considered a vocalization to be reliably assignable if the Mouse Probability Index (MPI) exceeded 0.95, where the MPI was defined as

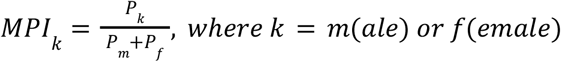

and *P*_*k*_ is the probability that the currently localized USV originated from the male or female, computed as 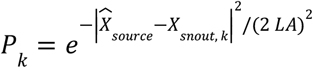, where 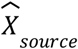 is the position estimate by SLIM, *X*_*snout*, *k*_ the position of the snout of animal *k* and *LA* the localization accuracy as defined above (normalization factors omitted, as they drop out in the *MPI*_*k*_). Hence, we assume a normal distribution of locations around the snout of each animal with a standard deviation given by the accuracy of localization via SLIM.

Similar to the absolute exclusion criterion by (Neunuebel et al., 2015), we also excluded USVs from analysis if they were localized >10 cm away from either mouse. These two criteria reduced the total set by ∼15% (22,220/26,363 USVs kept for Exp. 1; for Exp.2 with 3 microphones 4738/11,729 USVs were kept, i.e. ∼40%), which formed the set on which the subsequent analyses were based.

We investigated the influence of these selection criteria on the accuracy and the fraction of USVs kept for analysis in a simulation (Fig. 3, Supporting Data 3). The simulation was based on the distribution of inter-snout distances during the experiment (see Fig. 1D). We drew 10^5^ random samples {*d*_*i*_} _*i*=1…100000_ from the distance distribution and placed an emitter at the coordinates (0,0) and a receiver animal at (*d*_*i*_,0). Then, randomly drawn location estimation errors were added to the source animal in 2D, drawn from a normal distribution with a particular MAE. This procedure was repeated for MAEs ranging between 1 and 100 mm in 1 mm increments. The resulting location was then assigned to either the emitter or receiver animal based on proximity. The accuracy was quantified as the percentage of USVs assigned to the emitter (Fig. 3 SD3 A, light green). Results were also filtered with the selection criteria. Further, the simulation was repeated conditionally on particular interanimal distances (Fig. 3 SD3 B/C), which, when applying the above criteria, highlights the difference in accuracy for snout-snout interaction and for algorithms with different average MAEs.

USVs assigned to a single mouse were included in the subsequent analysis of vocalizations, in particular the relative spatial positions of the mice during USV production and the associated USV properties (Fig. 4 and 6). The relative spatial position of the receiver mouse relative to the emitter mouse was estimated in polar coordinates. The coordinate system was based on the snout of the emitter mouse (see Fig. 4A), with the line between the head centre and the snout pointing towards 0° (which was plotted pointing upwards in the plot). The vector pointing to the receiver mouse was rotated appropriately and converted to a polar representation. We assumed that the mice had no preference for relative vocalizations on the left/right to their snout and all vectors were thus mirrored to the right side for further analysis. The data points (2d vectors) were then binned using a polar histogram with evenly sized bins across angle and radius.

This resulted in a raw count histogram of relative positions during USV emission for male and female mice (Fig. 4B). As this histogram is biased by the general distribution of relative positions the animals took with respect to each other, it was normalized pointwise via the histogram of relative positions collected over all video frames (see Fig. 4A). In this way, we obtained the conditional relative spatial vocalization density for both sexes (Fig. 4C).

Further, we quantified the relative spatial distribution of various USV properties (see next section) by averaging the corresponding properties of all USVs contributing to a particular bin of the raw count histogram (Fig. 5). In the depiction, the hue indicates the average property, whereas the intensity (controlled via the transparency) indicates the normalized occurrence density. In this way, only intense colours indicate sufficient sampling of a bin to compute a meaningful average.

### USV Analysis

We used a range of techniques to estimate derived properties of each USV. First, we used the same set of automatically extracted acoustic and shape properties (see VocAnalyzer.m) as in (Ivanenko et al., 2020). In total, 17 scalar and 3 vectorial properties were estimated for each USV (see (Ivanenko et al., 2020) for details). Second, we performed nonlinear dimensionality reduction and nonlinear clustering in a range of configurations to assess the grouping of USVs (see `VocClassifier.m`).

The dimensionality reduction analysis was based on the fundamental frequency line of each USV, i.e. the sequence of frequency values of the fundamental frequency over time (see also (Ivanenko et al., 2020)), its derivative (each discretized at 1 ms for up to 100 ms, i.e. 100 dimensions each), USV duration, average directionality (ascending/descending in frequency), and the snout distance, which together constitute 203 parameters. We used the recently developed dimensionality reduction technique UMAP (McInnes et al., 2018), which is considered to provide better grouping than PCA while avoiding the variability in results associated with tSNE (van der Maaten and Hinton, 2008). The results of UMAP were stable and exhibited rich substructure (see Fig. 5). We used subsequent k-means clustering to group the vocalizations. Given the rich substructure, a large number of clusters was required to capture the subgroupings. However, because the clusters appeared to be connected, we considered the clustering mostly for visualization purposes.

We quantified the degree to which a property contributed to explaining the spatial structure of the USVs after dimensionality reduction by computing a measure of explained variance, 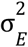, defined on the basis of local predictability. Concretely, we used nearest neighbor prediction (MATLAB function: knnsearch, with 5 nearest neighbors) to predict the entire dataset from its local congruency of a given property, e.g. mean frequency, duration, etc. This yielded a prediction error, here referred to as Local Prediction Error (LPE), simply as the RMSE (root mean squared error) distance between the data and the prediction. The LPE of the original data was then related to the residual error of permutations of the same values of the property on the given spatial structure to define the explained variance measure:

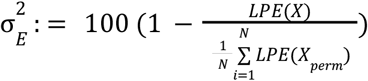

where the sum in the denominator runs over *N* = 100 random permutations and 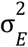 runs from 0 to 100%. We computed the significance as the number of permuted LPEs smaller than the original LPE divided by the number of permutations. The p-value, hence, could take values between 0 and 1 at a resolution of 0.01 (see Fig. 6B for a visualization of this analysis).

The dimensionality reduction was repeated using assumptions from previous publications (e.g. see (Holy and Guo, 2005; Neunuebel et al., 2015), namely (1) that USV duration is not important for USV classification (and all vocalizations can thus be stretched to a common length) and (2) that the mean frequency is not important for USV classification (and all USVs can thus be centred around 0 by subtracting their mean frequency). The results are shown in the two supporting data figures for Fig. 6.

### Statistical Analysis

To avoid distributional assumptions, all statistical tests were nonparametric, i.e. Wilcoxon rank sum test for two-group comparisons and Kruskal-Wallis for single factor analysis of variance. For the main statistical analysis in Fig. 5, we used a 3-way, nested ANOVA with sex, genetic variant and individual animal as predictors, where individual animals were nested inside the first two variables. Correlation is computed as Spearman’s rank-based correlation coefficient. Error bars represent standard errors of the mean (SEM) unless stated otherwise. All statistical analyses were performed in MATLAB v.2018b (The Mathworks, Natick) using functions from the Statistics Toolbox.

---

**Figure 3 - supporting data 1:**
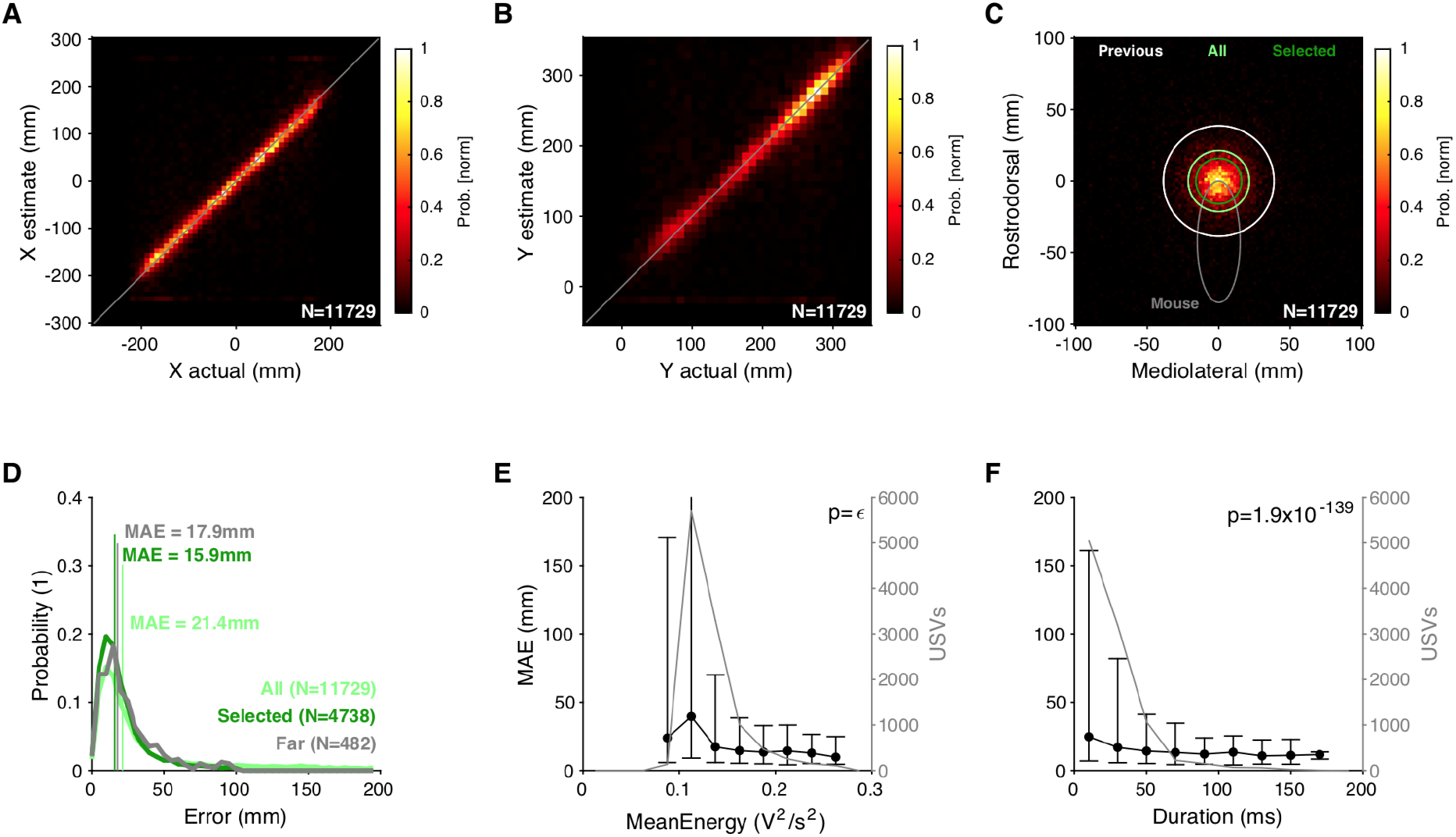
SLIM still provides high quality localization of vocalizations for 3 microphones. In Exp. 2, CBA/CaJ WT mice were recorded during social interaction using 3 microphones (located in an isosceles triangle nearly enclosing the platform in area, with the front two microphones in the same location as in Exp. 1 and the third one behind it in the middle), the minimal number of microphone necessary for SLIM to work. The results are comparable overall but less accurate than with 4 microphones (**A-C**), i.e. with an MAE = 21.4mm and 15.9 mm for all and the selected (see *Methods* for criteria) USVs, respectively. The percentage of reliably assignable USVs is substantially lower at 40.4%. (**D**). The dependence on mean energy (**E**) and duration (**F**) had a similar shape as for 4 microphones.

**Figure 3 - supporting data 2:**
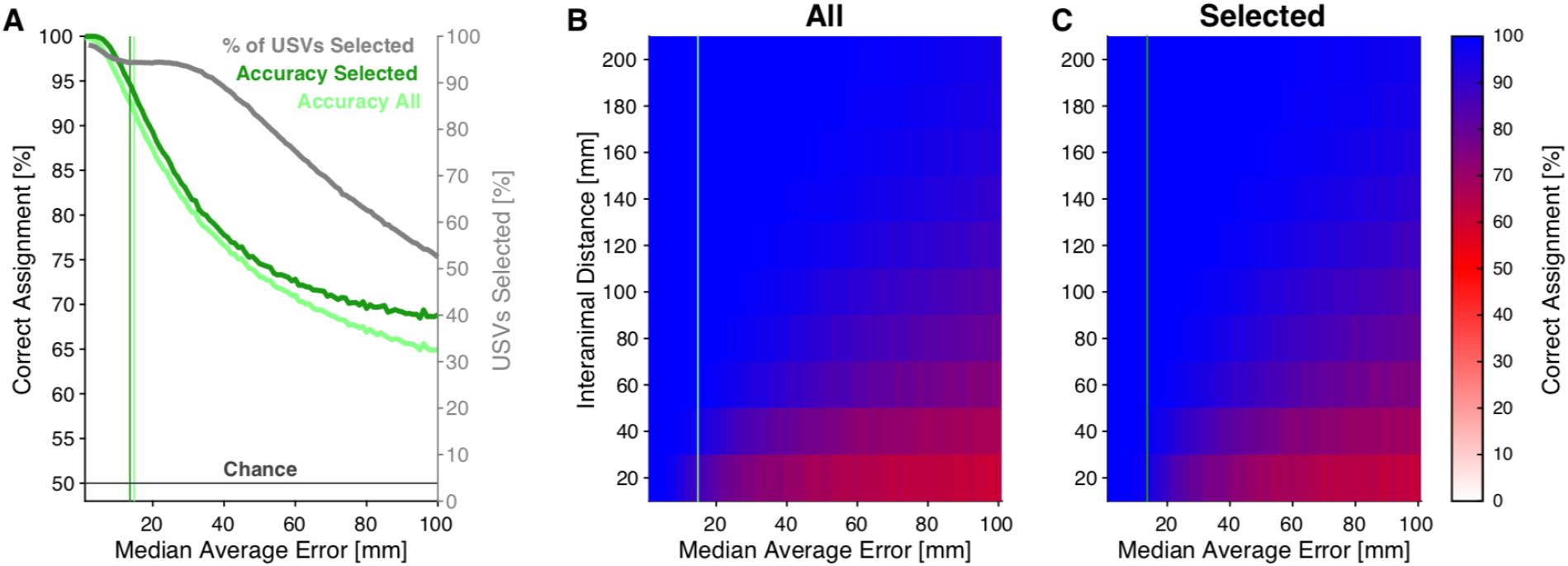
Influence of MAE and Selection Criterion on Accuracy of Assignment. We assessed the influence of our selection criteria on the resulting accuracy for different levels of median absolute errors by means of a simulation based on the empirical distance distribution of mice during interaction in our experiment (see Fig. 1D). **A** The accuracy decreased monotonically for increasing MAEs as expected. Applying our selection criteria (ii) and (iii) (see *Methods*), the accuracy increased substantially, with a widening margin for greater MAEs. The fraction of USVs that passed these criteria decreased strongly with MAE but remained high around the empirical MAE in our study (shown by light and dark green vertical lines, for All and Selected USVs, respectively). **B/C** In a second simulation we split up these results conditionally on interanimal distance, which demonstrated the expected result that for small interanimal distances the accuracy can be low for high MAEs. However, for the selected (reliably assignable) USVs and given the empirical MAE, the accuracy remains rather high at ∼90%, even for close snout-snout contact (range of up to 2cm).

**Figure 4 - supporting data 1:**
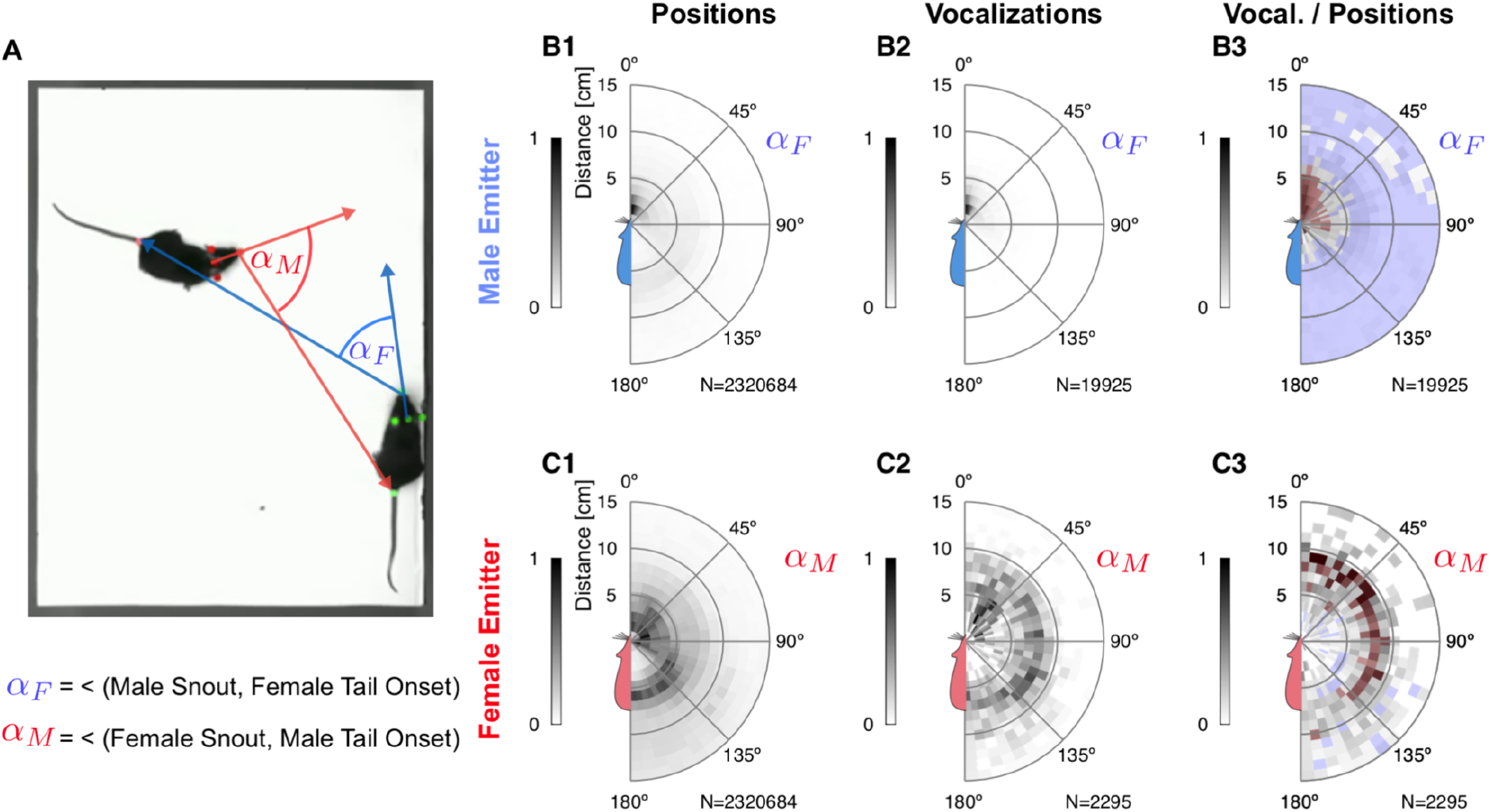
Relative emitter-snout and receiver-tail spatial densities during social interaction. The figure follows the same layout as Figure 4, with the receiver-head replaced by the receiver-tail-onset in the analysis as shown by the arrows in **A**. The males vocalized mainly around the tail-onset location of the female (**B3**), while the females vocalized mostly in snout-snout contexts, where the tail-onset of the male mouse is ∼6-10 cm away (**C3**); compare this to Fig. 4C3 as well.

**Figure 5 - supporting data 1:**
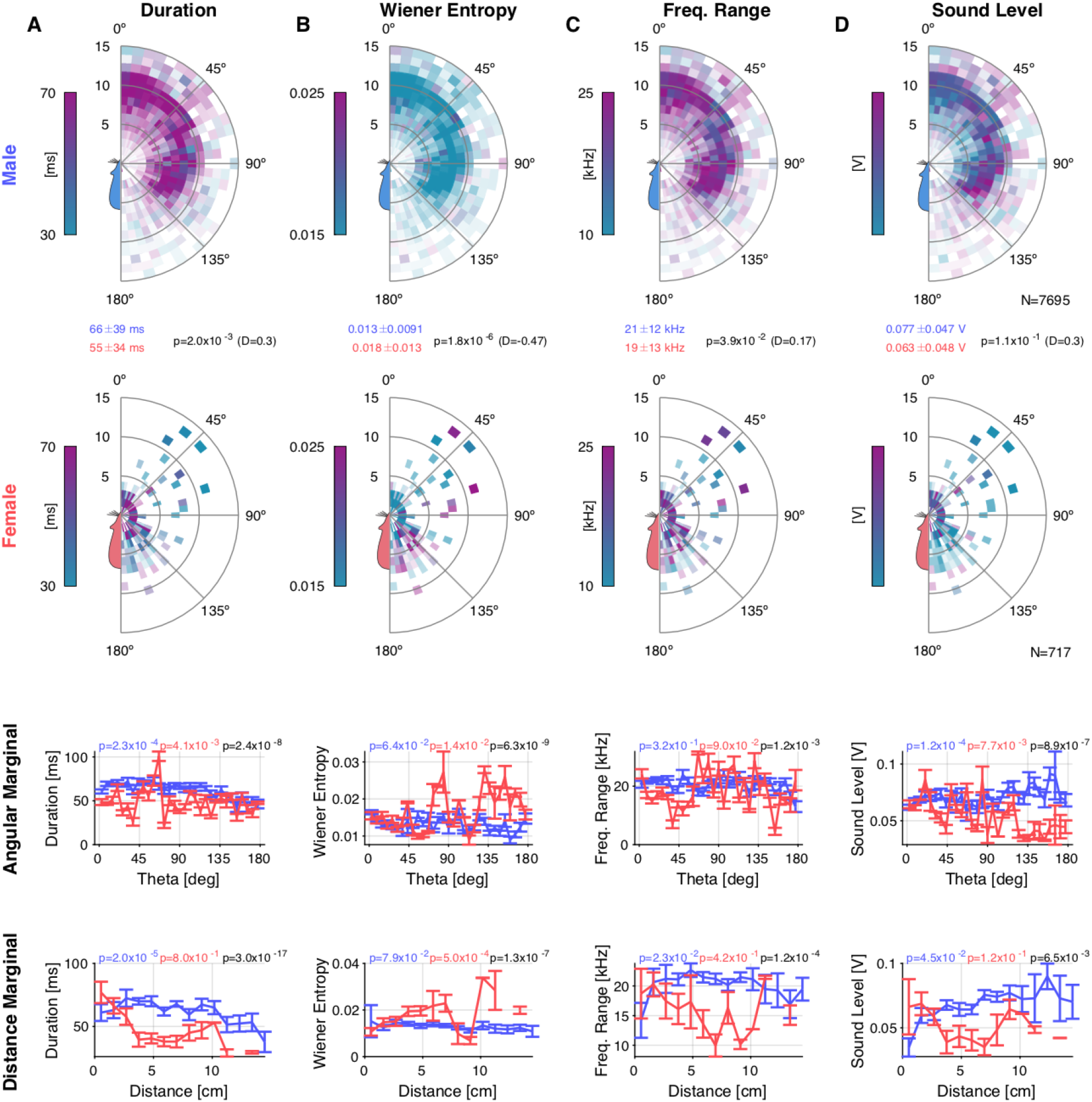
Figure 5: Properties of USV production depend on relative position and sex. **A** The duration of male USVs was substantially longer for males during snout-anogenital interaction than that of the duration of female USVs (comparison of top two rows). Male USV duration depended on both angle and distance (bottom rows), while female USV duration depended only weakly on these two factors. **B** The Wiener entropy of female USVs was greater than those of male USVs, in particular during snout-anogenital or snout-snout interactions where the male was behind the female. **C** The frequency range of male USVs was larger than that of female USVs, and they differed significantly as a function of distance and angle. **D** While there was no overall difference in sound level between male and female mice (see top), the spatial distributions show different patterns of dependence as a function of distance and angle, with male vocalization levels exceeding those of females for distances >5 cm and angles >100°, indicating snout-anogenital contact with the animals facing in opposite directions.

**Figure 6 - supporting data 1:**
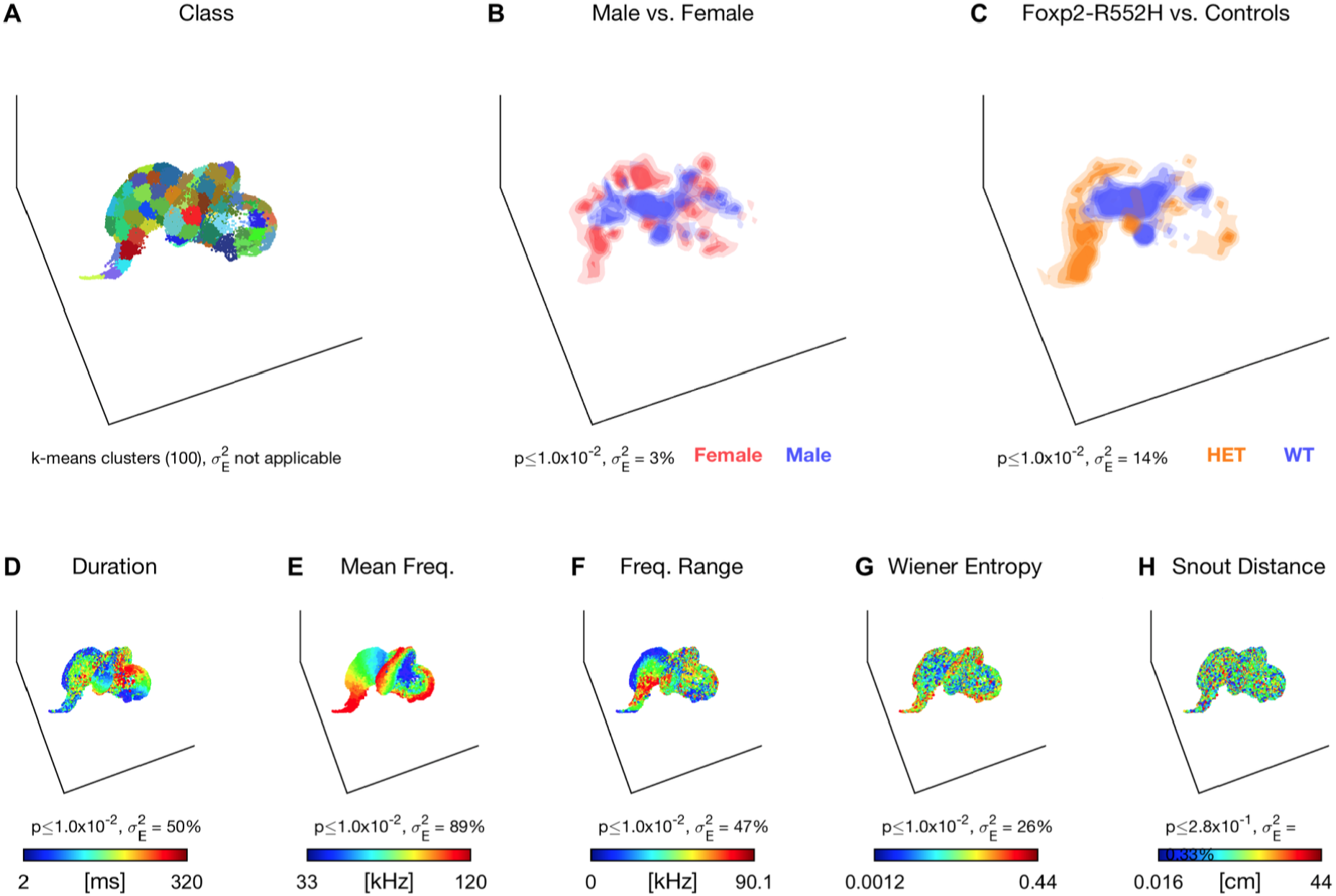
Same analysis as in Fig. 5, but with duration removed from the analysis by first stretching the USV to the same length as often done in other studies. The explained variance by the Foxp2-R552H variant dropped slightly, consistent with the differences in USV duration in Fig. 6 (N.B., the latter are spatially resolved, while these are partially averaged). The explained variance of the different genotypes increased to 14%, pointing to a variant-specific difference in the shape of the USVs. Note that duration, despite its removal, still had predictive value due to its correlation with other properties, e.g frequency range. Supplemental Movie 3 shows the same data revolving in 3D, resolving depth ambiguities.

**Figure 6 - supporting data 2:**
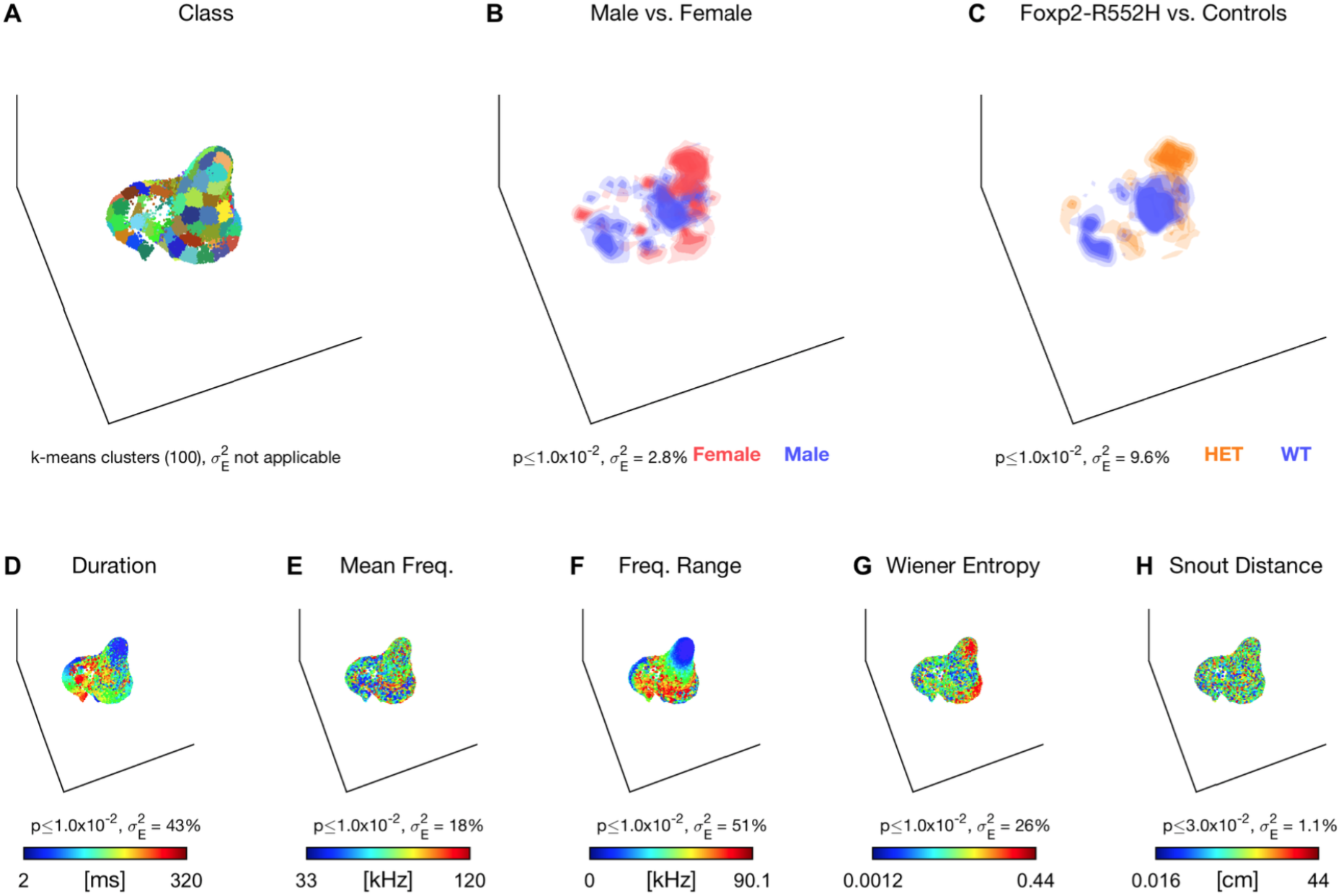
Same analysis as in Fig. 6, but now with duration and mean frequency removed (i.e. all vocalizations centred on the same frequency). As a consequence, the variance that duration and mean frequency explain ends up substantially lower while frequency range slightly increases. The explained variance by genotype is still low, indicating that frequency differences are contributing to the genotype differences in vocalization. Supplemental Movie 4 shows the same data revolving in 3D, resolving depth ambiguities.

## Notes

### Competing Interest Statement

The authors have declared no competing interest.

